# Neuron-astrocyte Coupling in Lateral Habenula Mediates Depressive-like Behaviors

**DOI:** 10.1101/2024.11.03.621722

**Authors:** Qianqian Xin, Junying Wang, Jinkun Zheng, Yi Tan, Xiaoning Jia, Zheyi Ni, Jiesi Feng, Zhaofa Wu, Yulong Li, Xiaoming Li, Huan Ma, Hailan Hu

## Abstract

The lateral habenular (LHb) neurons and astrocytes have been strongly implicated in depression etiology but it was not clear how the two dynamically interact during depression onset. Here, using multi-brain-region calcium photometry recording in freely-moving mice, we discover that stress induces a unique, bimodal neuronal response and a most rapid astrocytic response in the LHb. LHb astrocytic calcium requires the α_1A_-adrenergic receptor, and depends on a recurrent neural network between the LHb and locus coeruleus (LC). Through the gliotransmitter glutamate and ATP/Adenosine, LHb astrocytes mediate the second-wave activation of local LHb neurons as well as release of norepinephrine (NE). Activation or inhibition LHb astrocytic calcium signaling facilitates or prevents stress-induced depressive-like behaviors respectively. These results identify a stress-induced positive feedback loop in the LHb-LC axis, with astrocytes being a critical signaling relay. The identification of this prominent neuron-glia interaction may shed light on stress management and depression prevention.

## Introduction

Stress is a prominent risk factor for the development of depressive illness.^1^ Modern perspectives on the etiology of major depressive disorder (MDD) suggest that the molecular expression, cellular activities, and synaptic connections of specific brain regions are altered in response to external stressful stimuli.^2–4^ Recently, the lateral habenula (LHb), known as the brain’s ‘anti-reward center’ has been strongly implicated in the pathophysiology of major depression.^5–7^ LHb regulates essentially all neuromodulatory systems, including the serotonergic, dopaminergic, and noradrenergic systems.^7–9^ LHb neurons are activated by stress^10–13^ and are hyperactive under the depressive state.^14–19^ Although neurons have been the primary focus for depression etiology, recent studies have begun to implicate malfunctions of LHb astrocytes, including increased expression of Kir4.1 for potassium buffering^20^ and reduced expression of GLT1 for glutamate uptake.^21,22^ However, it has remained elusive how neurons and astrocytes in the LHb coordinate their activity in response to stress, and how their dynamic interaction contributes to depression etiology.

The synergy between neurons and astrocytes is increasingly recognized as crucial in brain physiology, animal behavior, and disease.^23–28^ Astrocytes communicate with neurons through chemically based dialogue.^29,30^ Astrocytes possess dense fine processes and express various types of G protein-coupled receptors (GPCRs), bestowing them to sense neuronal and synaptic activities with intracellular calcium transients.^31–35^ Astrocytic calcium levels can be elevated by neuromodulators, such as acetylcholine (ACh),^36–38^ dopamine,^32,39^ serotonin^40^ or norepinephrine (NE).^41–43^ In turn, astrocytes can regulate neuronal functions at diverse timescales through the uptake of extracellular ions and neurotransmitters^20,44–46^ or release of gliotransmitters.^27,30,47–52^ Gliotransmitters, including glutamate, gamma-aminobutyric acid (GABA), D-serine and ATP/Adenosine, can lead to synchronized neuronal excitation,^29,53–55^ enhanced temporal fidelity of firing,^56^ or fine-tuned synaptic transmission.^32,39,57–61^ However, in the context of stress response and depression, the bi-directional signaling of how neurons and astrocytes communicate with each other has remained much less studied.

In this study, we set out to apply fiber photometry in freely-moving mice to record the calcium activities of neurons and astrocytes in response to stress in multiple brain regions. We found that stress induced a most rapid astrocytic response and a unique, biphasic neuronal response in the LHb. Using cell-type-specific manipulations in combination with fiber photometry and two-photon imaging recordings, we identified a bidirectional neuron-astrocyte interaction in the LHb-LC circuit during stress response, and pinpointed a critical role of LHb astrocytes in the onset of depression.

## Results

### Foot-shock-stress induces most rapid calcium response in LHb astrocytes

We performed in vivo multi-site fiber photometry to record astrocytic calcium activities during foot-shock stress (FS, duration: 1 s; intensity: 1 mA; Figure 1A). We recorded a total of 6 brain regions known to be responsive to stress, including the medial prefrontal cortex (mPFC), hippocampus (HPC), lateral habenula (LHb), lateral hypothalamus (LH), striatum (STR) and basolateral amygdala (BLA). Adeno-associated viral (AAV) 2/5 expression of the glia-specific GfaABC1D promoter^62^ driving the genetically encoded calcium indicator GCaMP6f^63^ allowed the recording of calcium signals specifically in astrocytes (Figure 1B). Among the GCaMP6f positive cells, 91.9% were S100β (astrocytic marker) positive and 0.3% were NeuN (neuronal marker) positive (Figure 1C and S1A), suggesting a highly specific expression. Three brain regions were simultaneously recorded per animal and their calcium responses were aligned to the onset of FS to allow the comparison of signals across different brain regions (Figure 1D and 1E). While all six recorded regions showed increased astrocytic calcium activities in response to FS (Figure 1E), signals in the LHb peaked significantly earlier than those of other five regions (mean latency to peak 1.68 ± 0.08 s in LHb, 2.22 ± 0.12 s in LH, 3.17 ± 0.24 s in PFC, 3.37 ± 0.22 s in HPC, 3.45 ± 0.13 s in BLA, 4.98 ± 0.89 s in STR; LHb versus PFC, HPC, BLA, STR, p < 0.0001; LHb versus LH, p = 0.001; Mann-Whitney test; Figure 1D, 1E and 1G). The peak amplitude of FS-evoked LHb astrocytic calcium signals was also larger than those of other five regions (LHb versus LH, PFC, p = 0.01; LHb versus HPC, p = 0.005; LHb versus BLA, STR, p = 0.0002; Mann-Whitney test; Figure 1H). By analyzing the kinetics of calcium signals in mice with varying levels of peak fluorescence in the LHb, we found that the time to peak did not depend on the peak amplitude (Figure S1B), suggesting that the kinetics of calcium signals is not affected by signal intensity, but rather reflects intrinsic regional properties. Thus, among the multiple regions examined, the LHb astrocytes showed the most rapid response to FS, suggesting that it may play a unique role in processing stress-related information.

**Figure 1.**
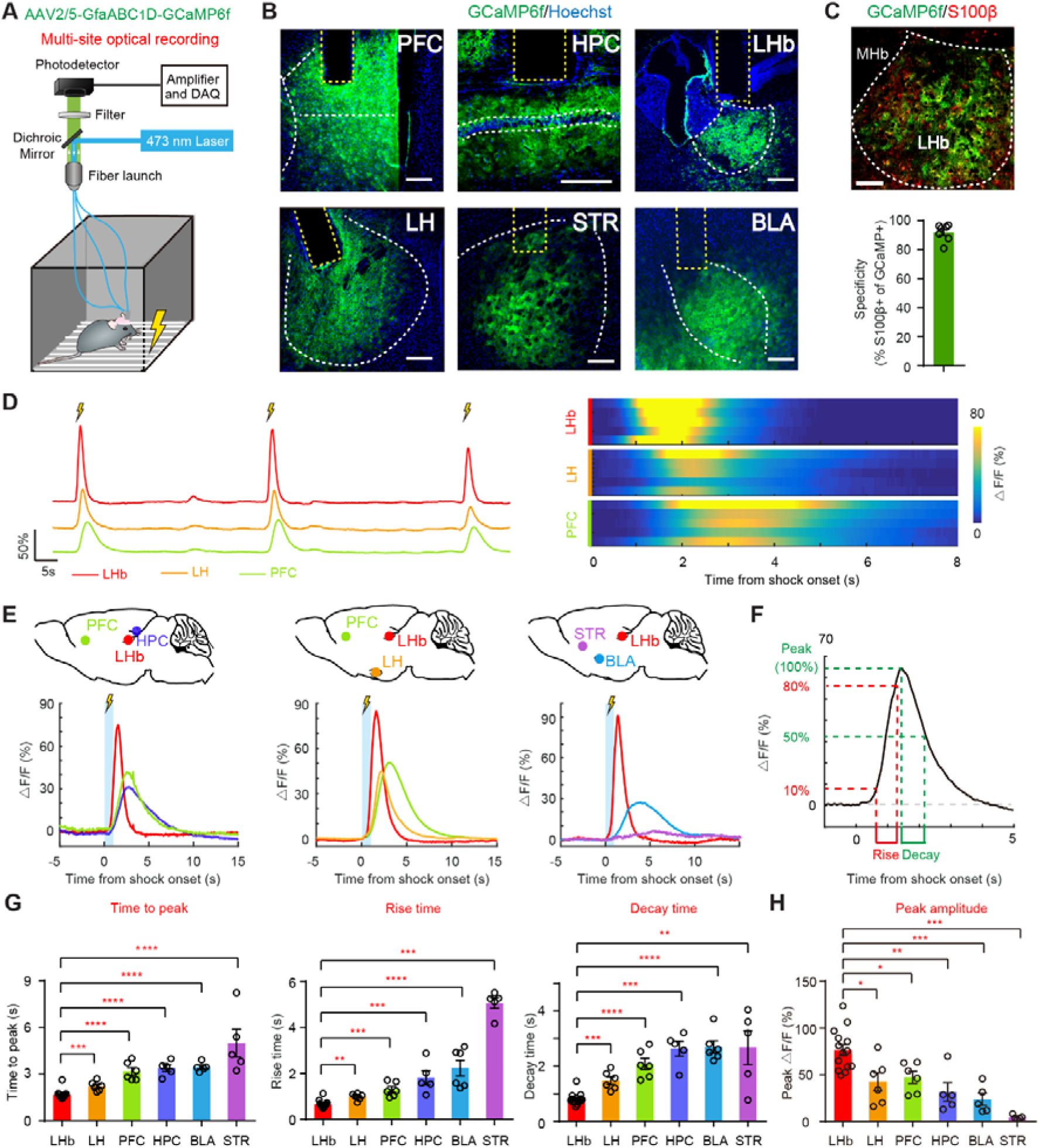
Foot-shock-stress Induces Most Rapid Calcium Response in LHb Astrocytes (A) Schematic illustrating viral construct for astrocyte-specific expression of GCaMP and multi-site fiber photometry setup for recording astrocytic calcium activities during FS. (B) Illustration of viral expression (stained with antibodies against GFP and Hoechst) and optic fiber placement (indicated by the yellow dotted line) in 6 central nervous system (CNS) regions, the boundary of which is outlined by the white dotted line. Green, GCaMP6f; blue, Hoechst. Scale bar, 200 μm. (C) GCaMP expression is highly specific in astrocytes. Top: representative image of LHb brain slices stained with antibodies against GFP (which indicates GCaMP positive cells, green) and S100β (astrocytic marker, red). Scale bar, 100 μm. Bottom: bar graph showing percentage of GCaMP-positive cells that are S100β-positive (n = 7 slices from 3 mice). (D) Representative traces (left) and heatmap (right) of delta F/F ratio of simultaneously recorded astrocytic calcium signals in LHb, LH and PFC over three continuous FSs. Scale bars, 50% delta F/F, 5s (left). (E) Alignment of FS-onset plots of calcium signal changes of simultaneously recorded multi-region astrocytes during FS. (F) Illustration of calculation of rise and decay time. (G) Average time to peak (left), rise time (middle) and decay time (right) of astrocytic calcium signals evoked by FS in 6 brain regions. Each circle represents one mouse. (H) Average peak amplitudes of astrocytic calcium signals evoked by FS in 6 brain regions. Each circle represents one mouse. *P < 0.05; **P < 0.01; ***P < 0.001; ****P < 0.0001; n.s., not significant. Data are represented as mean ± SEM.

### Foot-shock-stress induces biphasic calcium activities in LHb neurons

We next expressed GCaMP6s under the CaMKIIα promoter in the above six brain regions and performed calcium photometry recording in neurons (Figure 2A and S2A-S2E). Consistent with previous reports that LHb neurons encode aversive signals,^10,12^ LHb neuronal calcium signals were strongly activated in response to FS stress (Figure 2B, 2C, S3A and S3B). Interestingly, the FS-evoked calcium signals in LHb neurons first peaked at 0.9 ± 0.03 s (Figure 2D), and was followed by a second phase of activation with the inflection point at 2.54 ± 0.16 s (Figure 2C, 2D and S3C). The mean half-decay time of the first fast and the second slower phase was 0.83 ± 0.05 s and 22.4 ± 1.77 s respectively (Figure S3D). Although the second-phase signal has a smaller peak, due to its prolonged decay time, its area under the curve (AUC) is 3.46 ± 0.78 folds of that of the first phase (Figure 2D and S3E), suggesting that even more calcium entered during the second phase. Such FS-evoked bimodal neuronal calcium activation was not observed in the other five brain regions examined (Figure S2).

**Figure 2.**
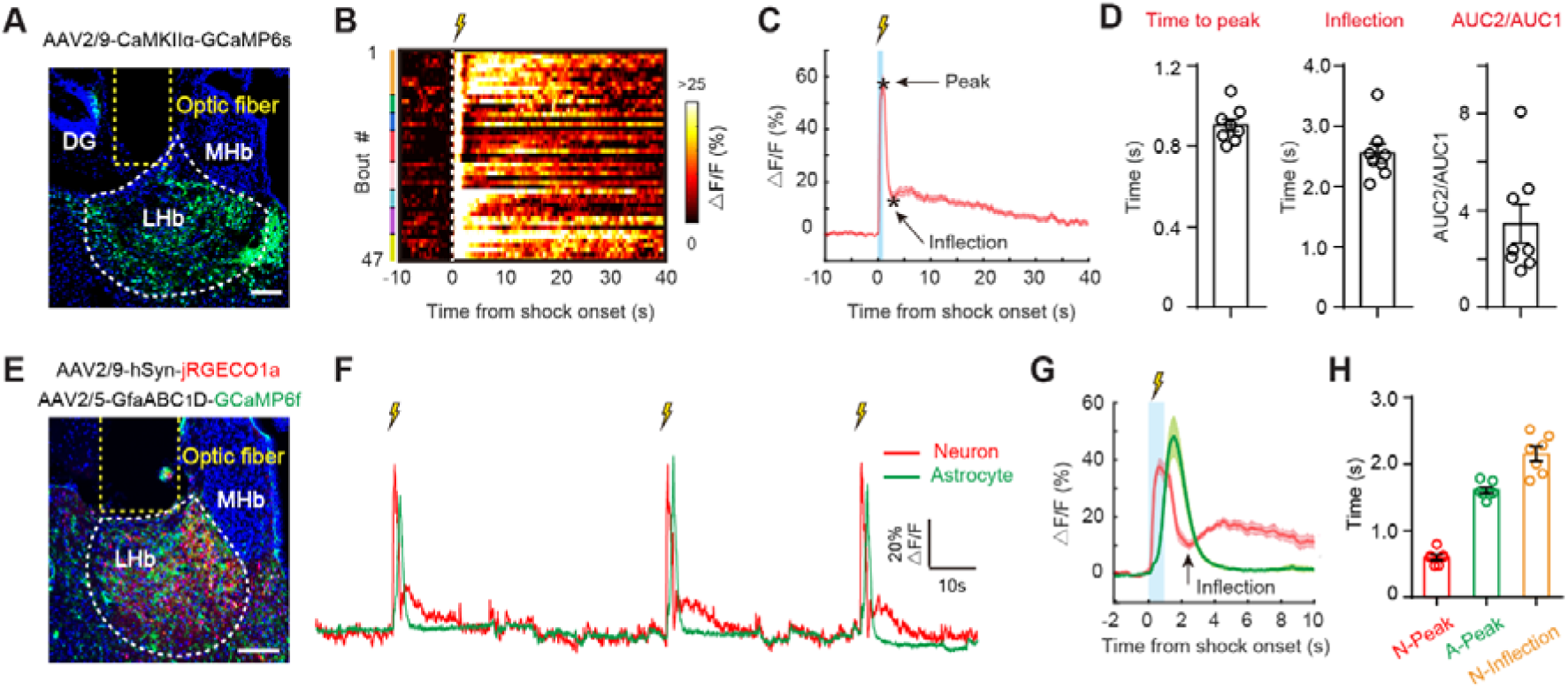
Foot-shock-stress Induces Biphasic Calcium Activities in LHb Neurons (A) Illustration of viral construct for neuron-specific expression of GCaMP, viral expression and optic fiber placement (indicated by the yellow dotted line) in the LHb, the boundary of which is outlined by the white dotted line. Green, GCaMP6s; blue, Hoechst. Scale bars, 100 μm. (B) Heatmap of delta F/F ratio of calcium signals from LHb neurons (n = 47 trials from 8 mice) aligned to onset of FS. Color bars on the left indicate different mice. (C) Average delta F/F ratio of calcium signals from LHb neurons aligned to onset of FS. Solid lines indicate mean and shaded areas indicate SEM. (D) Average time to peak (left), average time to inflection (middle) of LHb neuronal calcium signals from FS onset, and ratio of area under curve (AUC) of FS-evoked signals after and before the inflection point (right). AUC1: AUC of neuronal calcium signal in 0-inflection from FS onset. AUC2: AUC of neuronal calcium signal in inflection-10%peak. Each circle represents one mouse. (E) Illustration of dual-color neuronal expression of jRGECO1a and astrocytic expression of GCaMP6f and optic fiber placement (indicated by the yellow dotted line) in LHb. Scale bars, 100 μm. (F) Representative traces of simultaneously recorded neuronal and astrocytic calcium signals in LHb over three continuous FSs. Scale bars, 20% delta F/F, 10s. (G) Delta F/F ratio of calcium signals from LHb neurons and LHb astrocytes aligned to the onset of FS. Solid lines indicate mean and shaded areas indicate SEM. (H) Average time to peak of LHb neuronal (N-Peak, left) and astrocytic (A-Peak, middle) calcium signals and average time to inflection of LHb neuronal calcium signals from FS onset (N-inflection, right) in dual-color fiber photometry system. Each circle represents one mouse.

To better understand the relationship between stress-activated neuronal and astrocytic calcium activities in the LHb, we simultaneously recorded their activities by expressing the red calcium sensor jRGECO1a^64^ under the neuron-specific human synapsin (hSyn) promoter and green GCaMP6f under the GfaABC1D promoter (Figure 2E). Consistent with results from their separate recordings in Figure 1D-H and 2A-D, dual-color fiber photometry recording revealed that FS triggered sequential activation of LHb neurons and astrocytes, with the neuronal signals peaking at 0.64 ± 0.04 s and astrocytic signals peaking at 1.64 ± 0.05 s (Figure 2F-2H). It is also of interest to note that LHb astrocytic signals peaked between the first and second phase of LHb neuronal signals (Figure 2F-2H), suggesting a possibility that LHb astrocytes may be regulated by the first-phase neuronal activation, and contribute to the second-phase neuronal activity.

### Neuron-to-astrocyte crosstalk in the LHb during FS

In order to characterize the mechanism of LHb neuron-astrocyte interaction in stress processing, we first explored the potential crosstalk from neurons to astrocytes (Figure 3). We used AAV virus to express ChrimsonR^65^ in LHb neurons and GCaMP6f in LHb astrocytes. Subsequently, we recorded calcium activities in astrocytes following photostimulation of LHb neurons both *in vivo* and *in vitro* (Figure 3A and 3D). For the *in vivo* experiments, we tracked LHb astrocytic calcium activities using fiber photometry while optogenetically activating LHb neurons in freely moving mice (Figure 3A). For the *in vitro* experiments, we monitored LHb astrocytic calcium activities using two-photon imaging while optogenetically activating the LHb neurons in brain slices (Figure 3D). Surprisingly, while activation of LHb neurons robustly activated LHb astrocytic calcium signals in a frequency-dependent manner *in vivo* (Figure 3B and 3C), it failed to do so *in vitro* (Figure 3E and 3F). Even under stimulation frequency as high as 70 Hz, still no evoked astrocytic calcium signal was detected in brain slices (Figure 3F). We also coexpressed ChrimsonR and GCaMP6s in LHb neurons, and found that the same optogenetic stimulation protocol was able to strongly activate LHb neurons in brain slices (Figure S4A-S4D). These results suggest that first, activation of LHb neurons can lead to activation of LHb astrocytes *in vivo*; second, the lack of neuron-activated astrocytic calcium signals *in vitro* was not due to inefficient activation of LHb neurons, but more likely a lack of mediator of the neuron-astrocyte crosstalk in the brain slice setup.

**Figure 3.**
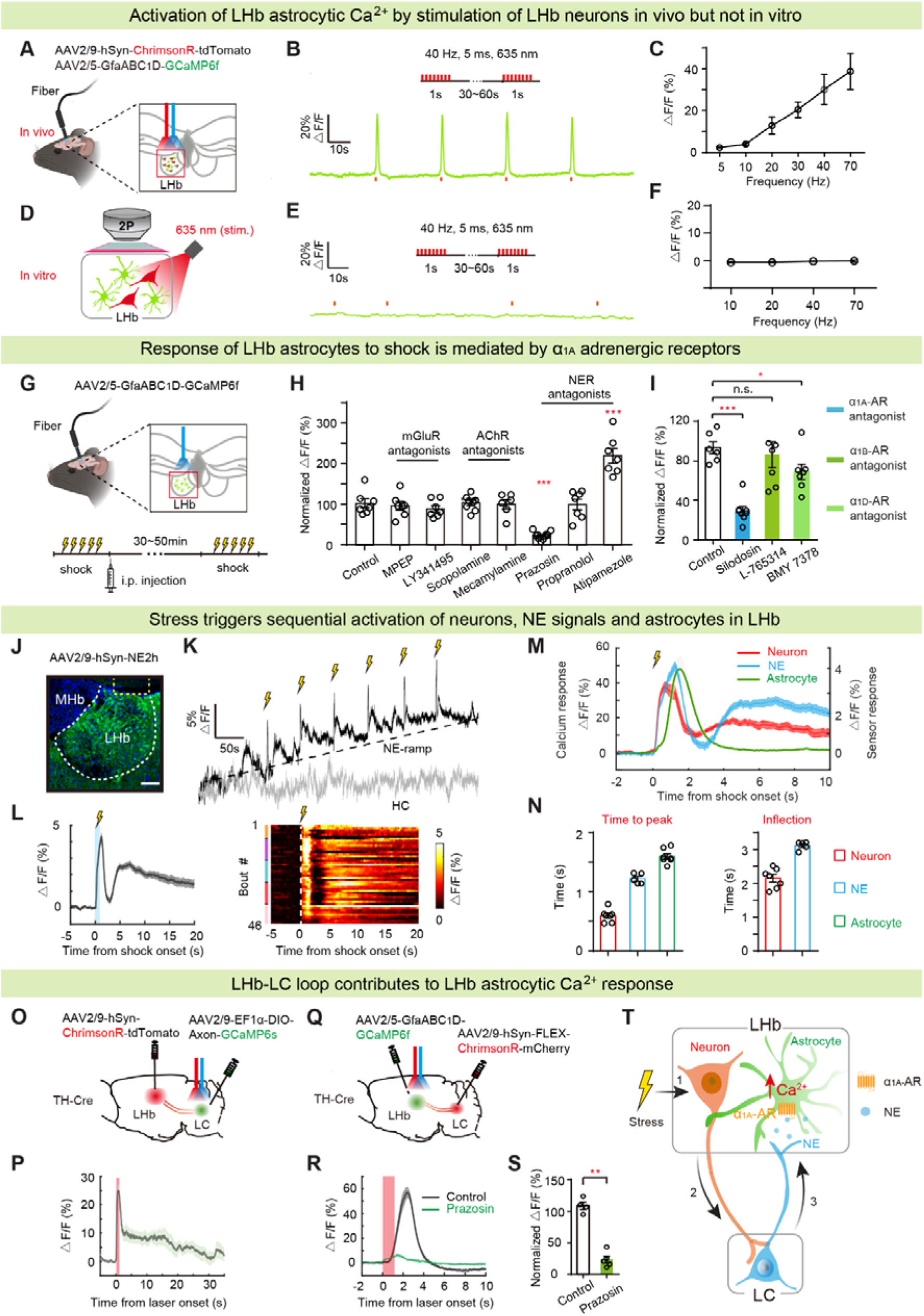
Neuron-to-Astrocyte Crosstalk in LHb during FS (A) Schematic illustrating optogenetic activation of ChrimsonR-expressing neurons and fiber photometry recording of nearby GCaMP6f-expressing astrocytes in LHb in vivo. (D) Schematic illustrating optogenetic activation of ChrimsonR-expressing neurons and two-photon imaging of nearby GCaMP6f-expressing astrocytes in LHb slices in vitro. (B, E) Astrocytic calcium responses to LHb neuronal ChrimsonR activation (1s, 40 Hz) in vivo (B) and in LHb slices (E). (C, F) Mean astrocytic calcium responses to varying stimulation frequencies in vivo (C) and in LHb slices (F). (G) Schematic of in vivo fiber photometry recording of LHb astrocytes in response to FS after i.p. injection of various receptor blockers. (H) Bar graph showing effects of MPEP (mGluR5 antagonist), LY341495 (mGluR2/3 antagonist), Scopolamine (mAChR antagonist), Mecamylamine (nAChR antagonist), Prazosin (α1-AR antagonist), Propranolol (β-AR antagonist) and Atipamezole (α2-AR antagonist) on FS-evoked calcium signals in LHb astrocytes. Each circle represents one mouse. (I) Bar graph showing effects of a1-AR subtype-selective antagonists, including Silodosin (α_1A_-AR antagonist), L-765314 (α_1B_-AR antagonist), BMY 7378 (α_1D_-AR antagonist) on FS-evoked calcium signals in LHb astrocytes. Each circle represents one mouse. (J-L) In vivo fiber photometry recording of NE sensor signals in LHb during FS. Viral expression (J, stained with antibodies against GFP and Hoechst) after injection of AAV2/9-hSyn-NE2h and optic fiber placement (indicated by the yellow dotted line) in LHb, the boundary of which is outlined by the white dotted line. Scale bar, 100 μm. Example trace (K) of LHb NE sensor signals during FS (black) and homecage (HC, grey). Scale bars, 5% delta F/F, 50s. Plots (L, left) of delta F/F ratio of NE sensor signals aligned to the onset of FS. Solid lines indicate mean and shaded areas indicate SEM. Heatmap (L, right) of delta F/F ratio of NE sensor signals in the LHb aligned to the onset of FS (n = 46 trials from 5 mice). Color bars on the left indicate different mice. (M) Plots of delta F/F ratio of calcium signals from LHb neurons, LHb astrocytes, and LHb-NE-sensor signals aligned to onset of FS stress. Solid lines indicate mean and shaded areas indicate SEM. Data from 2E-H and 3L. (N) Average time to peak of LHb neuronal calcium signals, LHb-NE-sensor signals and LHb astrocytic calcium signals from FS onset (left) and average time to inflection of LHb neuronal calcium signals and LHb-NE-sensor signals (right). Data from 2E-H and 3L. (O) Schematic illustrating optogenetic activation of LHb-LC terminals and fiber photometry recording of LC NE neurons in TH-Cre mice. (P) Calcium response of LC NE neurons to optogenetic activation of LHb-LC terminals (1s, 40 Hz). Plots of delta F/F ratio of calcium signal aligned to laser onset. Red line represents light-on. Solid lines indicate mean and shaded areas indicate SEM. (Q) Schematic illustrating optogenetic activation of LC^TH^-LHb terminals and fiber photometry recording of LHb astrocytes in TH-Cre mice. (R) Calcium response of LHb astrocytes to optogenetic activation of LC^TH^-LHb terminals (1s, 40Hz). Plots of averaged delta F/F ratio of astrocytic calcium signal induced by optogenetic activation of LC^TH^-LHb terminals after i.p. injection of saline (black) or Prazosin (green, α1-AR antagonist) aligned to laser onset. Red line represents light-on. Solid lines indicate mean and shaded areas indicate SEM. (S) Bar graph showing effects of Prazosin (α1-AR antagonist) on astrocytic calcium signals induced by optogenetic activation of LC^TH^-LHb terminals. Each circle represents one mouse. (T) Working model summarizing the neuron-to-astrocyte crosstalk in the LHb during FS. Stress activates LHb neurons, which recruit LC NE neurons to elevate LHb astrocytic calcium via NE-α_1A_ signaling. *P < 0.05; **P < 0.01; ***P < 0.001; ****P < 0.0001; n.s., not significant. Data are represented as mean ± SEM.

To search for the mediator of the neuron-astrocyte crosstalk in the LHb, we first looked for the astrocytic receptor(s) involved in this process *in vivo*. We measured astrocytic calcium photometry signals in response to FS while intraperitoneally (i.p.) injecting animals with blood-brain-barrier (BBB)-permeable inhibitors or antagonists of receptors for different neurotransmitters or neuromodulators (Figure 3G). We tested receptors for three classes of neurotransmitter/modulators, which were previously implicated in astrocytic Ca^2+^ signaling: the metabotropic glutamate receptors (mGluR),^31,66^ acetylcholine receptors (AChR) ^36,67,68^ and norepinephrine receptors (NER).^42,69,70^ While antagonists for mGluR5, mGluR2/3, muscarinic AChR (mAChR) or ionotropic AChR (nAChR), namely MPEP, LY341495, scopolamine and mecamylamine respectively, induced no change (p = 0.87 for MPEP and p = 0.52 for LY341495, p = 0.67 for scopolamine and p = 0.71 for mecamylamine, Mann-Whitney test), antagonists for the NE receptors caused strong impacts on FS-evoked LHb astrocytic Ca^2+^ (Figure 3H, S5A and S5B). In particular, α1-adrenergic receptors (α1-AR) antagonist prazosin largely suppressed FS-evoked LHb astrocytic calcium signals (p = 0.0001, Mann-Whitney test; Figure 3H and S5A). On the other hand, atipamezole, the antagonist of α2-adrenergic receptors (α2-AR), which are mostly expressed on presynaptic terminals serving autoinhibitory function,^71^ increased FS-evoked LHb astrocytic calcium signals (p = 0.0006, Mann-Whitney test; Figure 3H and S5B), possibly due to the release of autoinhibition. Among the α1-adrenergic receptors, there are three subtypes: α_1A_-, α_1B_- and α_1D_-adrenergic receptors (AR). By using their respective antagonists, we further delineated the primary contributor of FS-evoked LHb astrocytic calcium signals to be the α_1A_-AR (Figure 3I).

To further confirm that NE can activate LHb astrocytes through the α1-adrenergic receptors, we performed calcium imaging of astrocytes on LHb brain slices (Figure S6). Bath application of either NE or the α1-adrenergic receptor agonist phenylephrine (PE) increased astrocytic calcium signals in the presence of tetrodotoxin (TTX, which blocks all neuronal action potentials) (Figure S6A-S6E). The NE-elicited LHb astrocytic calcium signals were blocked by the α1-adrenergic receptor antagonist prazosin (Figure S6F and S6G). These data demonstrated that NE can activate astrocytic calcium signals through its action on astrocytic α1-adrenergic receptors in the LHb independent of local neural activity.

To detect whether NE is indeed released during FS in the LHb, we virally expressed in the LHb genetically encoded G-protein-coupled receptor activation-based NE (GRAB_NE_) sensor, NE2h (Figure 3J).^72^ Interestingly, in response to FS, NE signals exhibited both rapid phasic activity and slower ramping activity (Figure 3K and S7A-S7C). The rapid phasic increases in NE signals were time-locked to the delivery of FS, and similar as LHb neuronal Ca^2+^ signals, displayed bimodal distribution (Figure 3K and 3L). Over the course of 6 FSs, the LHb-NE signals kept ramping up and remained elevated for the duration of the 8 min session in the shock chamber (Figure 3K, S7A and S7C). Furthermore, alignment of LHb-neuron calcium, LHb-NE sensor and LHb-astrocyte calcium signals to the FS onset revealed a sequential activation of these three signals (Figure 3M and 3N).

The above data suggests that NE may be the potential link mediating the neuron to astrocyte crosstalk in the LHb. To further substantiate this hypothesis, we next explored the anatomical and functional connections between the LHb and LC, the hub for noradrenergic neurons and potential source of NE.^73–77^ First to look at the connection from the LHb to the LC, we injected AAV2/9 virus expressing hSyn promoter-driven ChrimsonR in the LHb, and AAV2/9 virus expressing Cre-dependent GCaMP6s into the LC of TH-Cre transgenic mice for NE-neuron-specific expression (Figure 3O and S8A). In these mice, optogenetic activation of LHb terminals in the LC evoked calcium response of LC-NE neurons (Figure 3P and S8B), suggesting that there is functional input from LHb neurons into the LC to activate NE neurons. Next, for the LC to LHb connection, we examined the samples from experiments in Figure 3O, and detected GCaMP signals from the TH^+^ LC neurons in the LHb (Figure S8A), verifying that anatomically there is a direct afferent input from the LC-NE neurons to the LHb region. Furthermore, optogenetic activation of LC-NE-neuron terminals in the LHb evoked calcium response of LHb astrocytes (Figure 3Q, 3R, S8C and S8D), which was blocked by α1-adrenergic receptor antagonist prazosin (p = 0.009, Mann-Whitney test; Figure 3S). Prazosin also abolished LHb astrocytic calcium signals evoked by activation of LHb neurons *in vivo* (p = 0.0095, Mann-Whitney test; Figure S8E-S8H). Furthermore, optogenetic inhibition of LHb neurons or chemogenetic inhibition of LC-NE neurons significantly decreased the calcium responses of LHb astrocytes evoked by FS stress (Figure S8I-S8P). Collectively, this set of data delineates a pathway, in which stress activates LHb neurons, in turn recruiting LC-NE neurons, which through the release of NE, stimulates the astrocytic calcium in the LHb (Figure 3T).

### Astrocyte-to-neuron crosstalk in the LHb

We next explored the signaling pathway mediating the potential crosstalk from LHb astrocytes to neurons. We first determined the effect of LHb astrocytic activation on LHb neuronal activities in freely moving mice (Figure 4A-4E). To mimic the activation of Gq-coupled α1-adrenergic receptor signaling,^78^ we expressed the Gq-coupled hM3Dq under the GfaABC1D promoter for chemogenetic activation of LHb astrocytes (Figure 4A). Upon injection of the ligand of hM3Dq clozapine, in vivo fiber photometry recording of the CaMKIIα-GCaMP6s-expressing LHb neurons revealed significantly elevated, dose-dependent, bulk calcium activity (Figure 4B and 4C). There was also an increase in the immunohistochemical signals of c-Fos, a cellular marker of cell activation, in LHb neurons (Figure 4D-4E).

**Figure 4.**
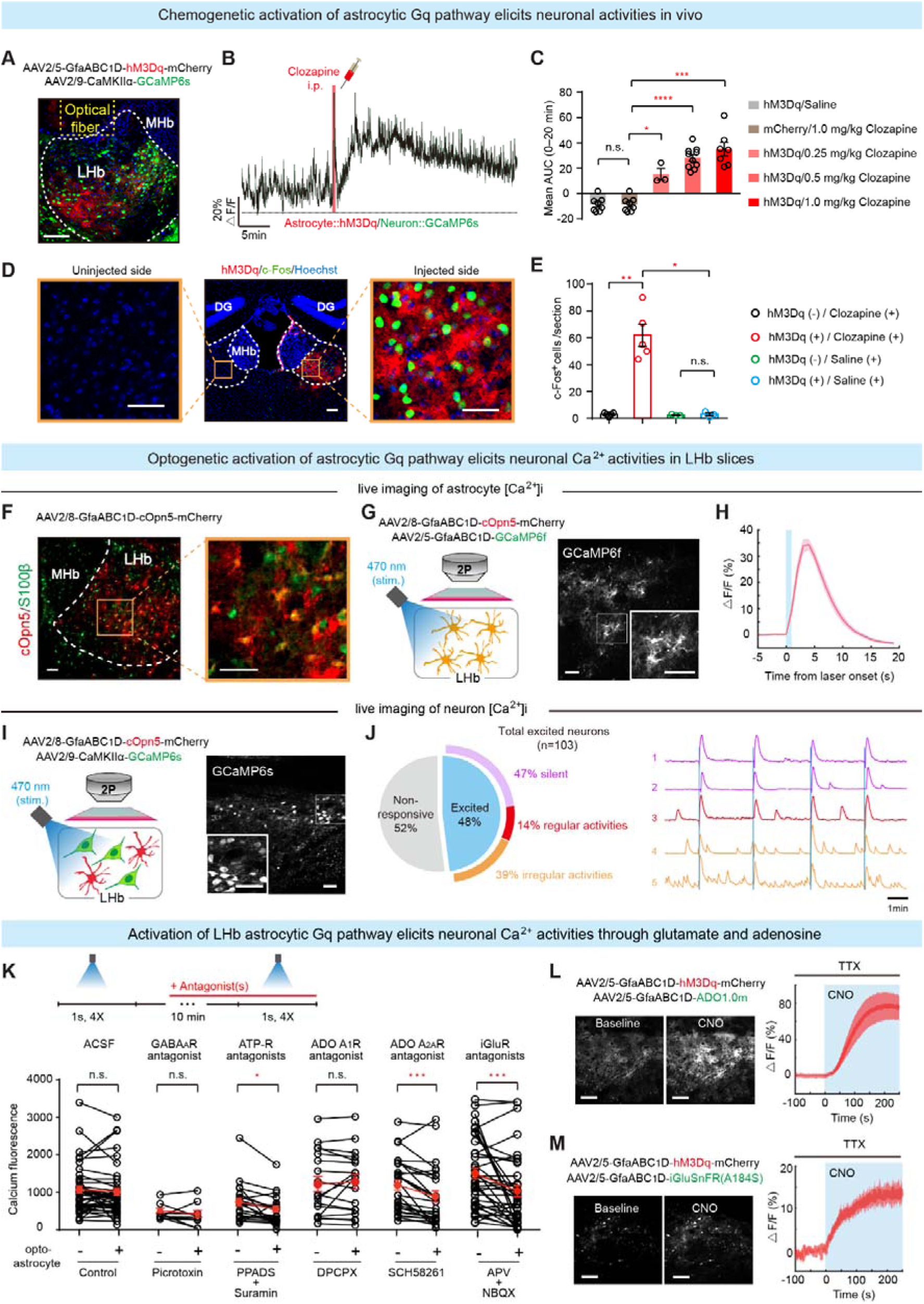
Astrocyte-to-Neuron Crosstalk in the LHb (A) Representative image showing viral expression of GCaMP6s in LHb neurons and hM3Dq in LHb astrocytes. Scale bar, 100 μm. (B-C) In vivo fiber photometry recording of LHb neuronal calcium in response to LHb astrocytic hM3Dq activation. Representative raw trace (B) and bar chart (C) showing mean AUC of LHb neuronal calcium signals following an intraperitoneal injection of saline or clozapine. Each circle represents one mouse. (D) Representative images of c-Fos IHC signals following unilateral astrocytic hM3Dq activation in LHb. Note c-Fos signals are only present in injected side. Green, c-Fos; blue, Hoechst. Scale bar, 40 μm. (E) Quantification of total c-Fos^+^ cells in LHb. (F) Representative image of LHb brain slices with viral expression of cOpn5 in astrocytes. Scale bar, 40 μm. (G) Schematic (left) and representative two-photon image (right) illustrating optogenetic activation of cOpn5-expressing astrocytes while recording two-photon images of LHb astrocytic calcium responses. Scale bar: 50 μm. (H) Plots of delta F/F ratio of calcium signals in LHb astrocytes aligned to laser onset. Solid lines indicate mean and shaded areas indicate SEM. (I) Schematic (left) and representative two-photon image (right) illustrating optogenetic activation of cOpn5-expressing astrocytes while recording two-photon images of LHb neuronal calcium responses. Scale bar: 50 μm. (J) Pie chart illustrating percent abundance of excited neurons by astrocytic cOpn5 activation (left, n = 4 slices from 4 mice). Colored outer circle indicates percentage of three types (according to baseline activity) among astrocytic-hM3Dq-excited neurons. Representative raw calcium traces from the three types of astrocytic cOpn5-excited neurons (right). Blue line represents light-on. (K) Bar graph showing effects of ACSF (control), picrotoxin (GABA_A_ receptor antagonist), PPADS and suramin (ATP-R antagonists, ATP receptor antagonists), DPCPX (ADO A_1_R antagonists, adenosine A1 receptor antagonist), SCH58261 (ADO A_2A_R antagonist, adenosine A_2A_ receptor antagonist) and APV and NBQX (iGlu receptor antagonists) on astrocytic cOpn5-evoked neuronal calcium signals in LHb slices. Each circle represents one neuron. (L, M) Illustration of AAVs used for expressing hM3Dq and ADO1.0m (L) or iGluSnFR (M) in LHb astrocytes (left top), fluorescence images (left bottom) and traces (right) illustrating astrocyte-induced adenosine (L) or glutamate (M) release in LHb slice. Scale bar: 100 μm. (Experiments were performed in TTX). *P < 0.05; **P < 0.01; ***P < 0.001; ****P < 0.0001; NS, not significant. Data are represented as mean ± SEM.

In LHb brain slices, calcium imaging recording also revealed successful activation of astrocytes and neurons upon chemogenetic activation of astrocytes (Figure S9A-S9F). In particular, 55% of the recorded neurons (148 of 267) showed significantly increased calcium signals upon CNO application (Figure S9F). Among these, 38% were silent, 55% and 7% showed irregular and regular baseline activity respectively before the CNO application (Figure S9F). To achieve astrocytic activation in a more transient and temporally controlled manner, we then tried optogenetic activation by expressing chicken opsin 5 (cOpn5), a light-sensitive Gq-coupled GPCR that can trigger a blue-light-induced elevation in intracellular calcium,^79^ in the LHb astrocytes (Figure 4F). Light stimulation (1 s, 473 nm) significantly evoked calcium activities in LHb astrocytes (Figure 4G and 4H). And the signals decayed much faster than the CNO-evoked ones (7.84 ± 2.25 s vs. 62.23 ± 23.92 s, Figure 4H and S9C), closer to the FS-evoked physiological condition (Figure 1G). Using this approach, we found that upon optogenetic activation of LHb astrocytes, 48% (103 of 216) recorded LHb neurons were excited (Figure 4I, 4J and S10A-S10B). Among these, 47% were silent, 39% and 14% showed irregular and regular baseline activity respectively before the stimulation (Figure 4J). Overall, both chemogenetic and optogenetic activation of astrocyte-Gq pathway significantly increased the activities of LHb neurons.

While the optogenetic activation of astrocyte-Gq may be less efficient than the chemogenetic method, it caused a more physiological change (in terms of signal duration) that allowed us to identify the gliotransmitter(s) mediating the LHb astrocyte-neuron crosstalk. Calcium signals in astrocytes are known to facilitate the release of various gliotransmitters, including glutamate,^80–82^ GABA^83,84^, ATP or its immediate degradation product adenosine.^61,85,86^ Therefore we measured LHb neuronal calcium activities triggered by optogenetic stimulation of LHb astrocytes in brain slices in the presence of antagonists for these gliotransmitter receptors (Figure 4K). While antagonists for the GABA_A_ (picrotoxin) and adenosine A_1_ (A_1_R, DPCPX) receptors did not show any effect (Figure 4K), those blocking adenosine A_2A_ (A_2A_R, SCH58261, Figure 4K and S10D) or ionotropic glutamate receptors (iGluR, APV+NBQX, Figure 4K and S10E) significantly suppressed astrocyte-stimulated neuronal calcium responses (34.8% reduction and p = 0.0002 for SCH58261; 32% reduction and p = 0.0008 for APV+NBQX; paired t test, Figure 4K). 52% and 44% of astrocyte-excited neurons exhibited significantly reduced calcium activities in response to adenosine A_2A_R and iGluR blockade respectively (Figure S10D and S10E). On the other hand, blockade of ATP receptors (ATP-R, PPADS + suramin) had a significant but much smaller effect on astrocyte-stimulated neuronal calcium responses (10% reduction and p = 0.012; paired t test, Figure 4K). Using genetically-encoded sensors of adenosine (ADO1.0m)^87^ or glutamate (iGluSnFR),^88^ we found that activation of astrocyte-Gq pathway (CNO, 1 μM, with TTX to exclude indirect release from neuronal activation) significantly increased the level of adenosine (Figure 4L) and glutamate (Figure 4M). These results suggest that activation of LHb astrocytes can lead to activation of LHb neurons through the release of gliotransmitters ATP/Adenosine and glutamate.

### LHb astrocytes account for the second-phase of LHb neural activity and NE release

We next investigated the functional impact of LHb astrocytic activities on the FS-evoked LHb neuronal activity *in vivo* (Figure 5). We employed GfaABC1D promoter to drive the expression of either a short hairpin RNA (shRNA) targeting intracellular IP3R2 for its knockdown (Figure 5A), or a 122-residue inhibitory peptide from β-adrenergic receptor kinase1 (iβARK, Figure 5K) to specifically block astrocytic calcium signaling. These approaches have been developed and validated to attenuate Gq-GPCR-mediated calcium signaling in astrocytes.^89,90^ Expression of these two viral constructs in the LHb significantly reduced FS-evoked calcium in LHb astrocytes by 68% and 51% respectively (p = 0.0002 for IP3R2-shRNA group, unpaired t test; p = 0.008 for iβARK group, Mann-Whitney test; Figure S11A-S11F). To examine the contribution of LHb astrocytes to neuronal calcium activities during FS stress, we then injected either one of these two viruses into the bilateral LHb and expressed GCaMP6s under CaMKIIα promoter in the unilateral LHb, and installed optic fibers above the LHb for in vivo photometry recording (Figure 5B, 5L, S11G and S11H). Notably, the second-phase LHb neuronal activity evoked by FS was largely eliminated by expression of IP3R2-shRNA (Figure 5C and 5D) or iβARK (Figure 5M and 5N) in LHb astrocytes. For the first-phase FS-evoked neuronal calcium signals, the time to peak, peak amplitude and AUC from 0 to 2.5s (AUC1) were not changed (Figure 5E-5G and 5O-5Q). However, for the second-phase FS-evoked neuronal calcium signals, the AUC from 2.5 to 40s (AUC2) was greatly reduced in mice expression IP3R2-shRNA compared with control (p = 0.005, Mann-Whitney test; Figure 5H and 5I). Consequently, the total AUC from the two phases was reduced by 55.7 % (p = 0.005, Mann-Whitney test; Figure 5J). Similar effects were observed in mice infected with iβARK (Figure 5R-5T). These results suggest that LHb astrocytic calcium activity are required for the second-phase LHb neuronal activity in response to stress.

**Figure 5.**
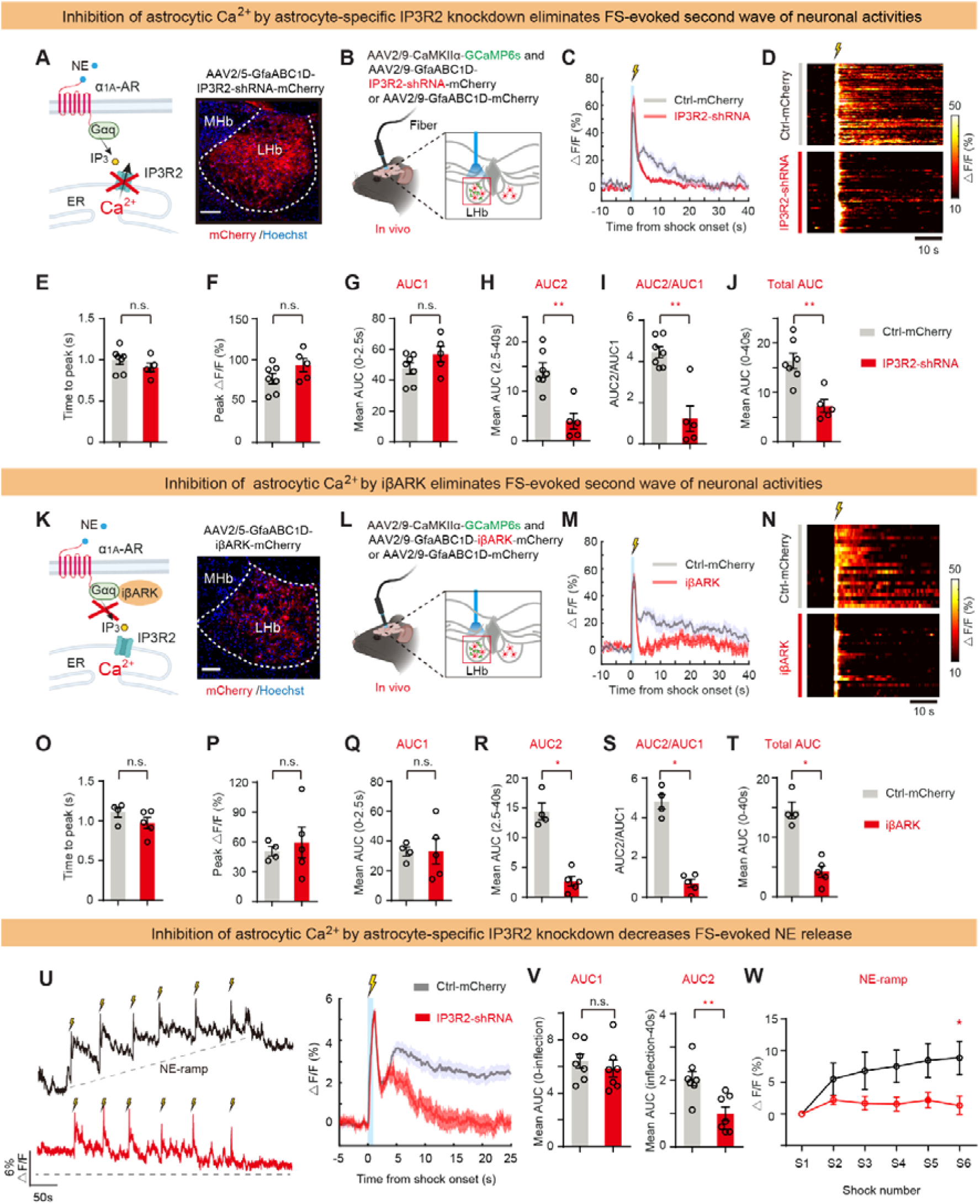
LHb Astrocytic Ca^2+^ Signaling Is Required for the Second-phase Neuronal Activities and NE Release (A) Left: illustration of blockade of astrocytic calcium signaling by IP3R2-shRNA. Right: viral expression of IP3R2-shRNA in astrocytes. Scale bar, 100 μm. (B) Schematic illustrating the viral expression of astrocyte-specific IP3R2-shRNA or Ctrl-mCherry in bilateral LHb and neuron-GCaMP6s in unilateral LHb, and fiber photometry recording of neurons in LHb. (C, D) Plots of averaged delta F/F ratio (C) and heatmap (D) of LHb neuronal calcium signals aligned to the FS onset in mice expressing Ctrl-mCherry (grey, n = 69 trials from 7 mice) or IP3R2-shRNA (red, n = 50 trials from 5 mice) in LHb astrocytes. (E-J) Quantification of Time to peak (E), Peak amplitude (F), AUC1 in 0-2.5s (G), AUC2 in 2.5-40s (H), AUC ratio (I) and total AUC in 0-40s (J) of FS-evoked LHb neuronal calcium signals in mice expressing Ctrl-mCherry or IP3R2-shRNA in LHb astrocytes. Each circle represents one mouse. (K) Left: illustration of blockade of astrocytic calcium signaling by iβARK. Right: viral expression of iβARK in astrocytes. Scale bar, 100 μm. (L) Schematic illustrating the viral expression of astrocyte-specific iβARK or Ctrl-mCherry in bilateral LHb and neuron-GCaMP6s in unilateral LHb, and fiber photometry recording of neurons in LHb. (M, N) Plots of averaged delta F/F ratio (M) and heatmap (N) of LHb neuronal calcium signals aligned to the FS onset in mice expressing Ctrl-mCherry (grey, n = 22 trials from 4 mice) or iβARK (red, n = 30 trials from 5 mice) in LHb astrocytes. (O-T) Quantification of Time to peak (O), Peak amplitude (P), AUC1 in 0-2.5s (Q), AUC2 in 2.5-40s (R), AUC ratio (S) and total AUC in 0-40s (T) of FS-evoked LHb neuronal calcium signals in mice expressing Ctrl-mCherry or iβARK in LHb astrocytes. Each circle represents one mouse. (U) Example trace (left) and plots of averaged delta F/F ratio (right) of LHb NE signals during FS in mice expressing Ctrl-mCherry (grey, n = 42 trials from 7 mice) or IP3R2-shRNA (red, n = 42 trials from 7 mice) in LHb astrocytes. (V, W) AUC1 in 0-inflection and AUC2 in inflection-40s of LHb NE signals aligned to FS onset (V) and basal (tonic) delta F/F ratio of LHb-NE signals in response to FS (W) in mice expressing Ctrl-mCherry (grey) or IP3R2-shRNA (red) in LHb astrocytes. *P < 0.05; **P < 0.01; ***P < 0.001; ****P < 0.0001; n.s., not significant. Data are represented as mean ± SEM.

Given the identification of the stress-induced positive feedback interactions between LHb and LC-NE neurons (Figure 3P and 3R) and between LHb neurons and astrocytes (Figure 3A-3C and 4A-4C), we next sought to examine whether LHb astrocytes can regulate NE release in the LHb. Following FS, both neuronal calcium and NE-sensor signals in the LHb showed a biphasic pattern, and the second peak of NE signals followed the second peak of neuronal calcium signals, both following the peak of astrocytic signals (Figure 3M and 3N). Such temporal pattern, combined with results in Figure 5A-T, suggest that the reactivation of LHb neurons by local astrocytes may lead to subsequent reactivation of LC-NE neuron and NE release. To test this possibility, we simultaneously expressed IP3R2-shRNA and NE sensor in the LHb and performed photometry recording during FS stress. Expression of IP3R2-shRNA significantly reduced the second-phase NE signals (Figure 5U and 5V). While the AUC of first-phase FS-evoked NE signals showed no difference, the AUC of second-phase FS-evoked NE signals was largely reduced in mice expressing IP3R2-shRNA compared with control (p = 0.004, Mann-Whitney test; Figure 5V). Additionally, the ramp-up of NE signals over multiple FSs was also inhibited (p = 0.026, Mann-Whitney test; Figure 5U and 5W). These results suggest that LHb astrocytes do not affect the first-phase, but strongly contribute to the second-phase NE release in response to stress.

### Activation of LHb astrocytes facilitates depression-like behaviors

We next explored the functional consequence of LHb-astrocytic activation on behavioral response to stress (Figure 6). We first tried to activate LHb astrocytes, by expressing GfaABC_1_D-hM3Dq in the LHb (Figure 6D), in a subthreshold stress protocol (1 mA, 1 s, 6◊ FS delivered within 6 min; Figure 6A). This protocol by itself did not induce depression-like phenotypes (Figure 6B). However, when LHb astrocytes were chemogenetically activated (i.p., DCZ, deschloroclozapine, 0.5 mg kg^-1^) before the first FS, mice developed depression-like phenotypes after this subthreshold protocol (Figure 6C-6E). Compared with the mCherry-expressing group, mice expressing hM3Dq in LHb-astrocytes were more immobile in the forced swim test (FST), which models behavioral despair, and showed less preference for sucrose water in the sucrose preference test (SPT), which models anhedonia (Figures 6E).

**Figure 6.**
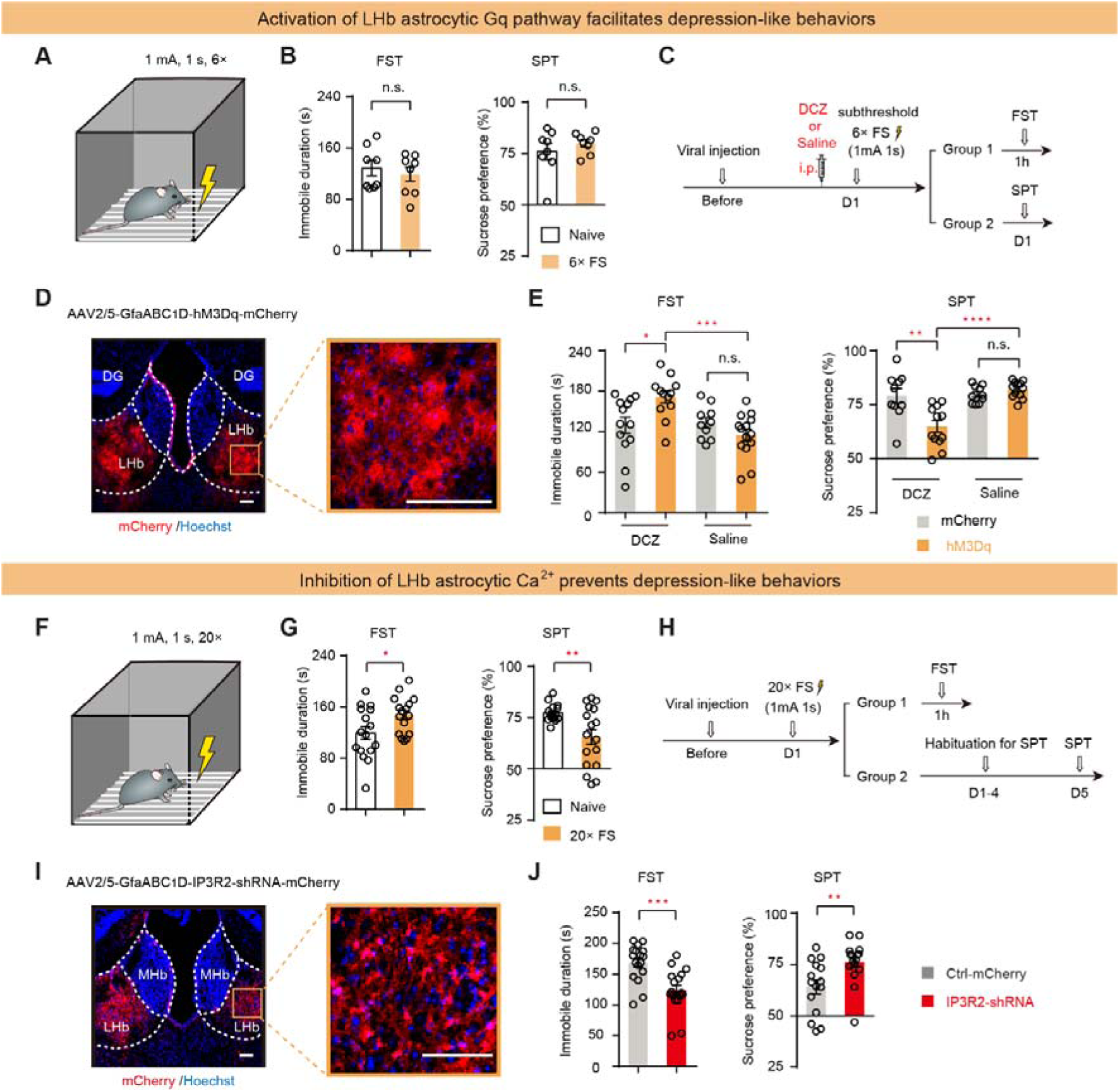
LHb Astrocytes Regulate Stress-induced Depression-like Behaviors (A) Cartoon illustration of a subthreshold depression-induction protocol with 6◊FS. (B) Subthreshold 6×-FS protocol did not induce depression-like behaviors in FST (n =8, 8 mice for naive and FS group, respectively) and SPT (n =8, 8 mice for naive and FS group, respectively). (C) Experimental paradigm for activating Gq pathway of LHb astrocytes during the subthreshold depression-induction protocol. (D) Illustration of viral expression of hM3Dq in LHb astrocytes. Scale bar, 100 μm. (E) Depression-like behaviors in FST (n =13, 12, 10, 14 mice for Ctrl-mCherry/DCZ, hM3Dq/DCZ, Ctrl-mCherry/saline and hM3Dq/Saline group, respectively) and SPT (n =10, 12, 10, 12 mice for Ctrl-mCherry/DCZ, hM3Dq/DCZ, Ctrl-mCherry/saline and hM3Dq/Saline group, respectively) of mice expressing Ctrl-mCherry (grey) or hM3Dq (blue) in LHb astrocytes. (F) Cartoon illustration of a depression-induction protocol with 20◊FS. (G) 20×-FS protocol induced depression-like behaviors in FST (n =16, 16 mice for naive and FS group, respectively) and SPT (n =15, 17 mice for naive and FS group, respectively). (H) Experimental paradigm for inhibiting calcium signaling of LHb astrocytes during the depression-induction protocol. (I) Illustration of viral expression of IP3R2-shRNA in LHb astrocytes. Scale bar, 100 μm (J) Depression-like behaviors in FST (n =17, 17 mice for Ctrl-mCherry and IP3R2-shRNA group, respectively) and SPT (n =15, 16 mice for Ctrl-mCherry and IP3R2-shRNA group, respectively) of mice expressing Ctrl-mCherry (grey) or IP3R2-shRNA (red) in LHb astrocytes. *P < 0.05; **P < 0.01; ***P < 0.001; ****P < 0.0001; n.s., not significant. Data are represented as mean ± SEM.

### Inhibition of LHb astrocytes prevents stress-induced depression-like behaviors

We next tested whether LHb-astrocytic activity is necessary for stress-induced depression. We tried to inactivate LHb astrocytes by expressing the IP3R-shRNA under the GfaABC1D promoter (Figure 6I). Mice were then exposed to a series of inescapable and unpredictable FS stress (1 mA, 1 s, 20◊FS delivered within 20 min; Figure 6F), which was shown to induce depression-like behaviors (Figure 6G).^16,91^ Compared with mice expressing the control shRNA, inhibition of LHb astrocytes with IP3R2-shRNA caused a pronounced reduction in depressive-like phenotypes (Figure 6H-6J), reducing immobility in the FST (p = 0.0003, unpaired t-test; Figure 6J) and increasing sucrose preference in the SPT (p = 0.0049, Mann-Whitney test; Figure 6J).

## Discussion

Among the six stress-related brain regions examined (mPFC, HPC, LHb, LH, STR and BLA), we found that LHb astrocytes and neurons exhibit the most rapid and unique, biphasic calcium pattern respectively in response to stress. Following FS, there is a sequential activation of neurons, release of NE from the LC, followed by activation of astrocytes in the LHb (Figure 3M and 3N). Manipulation experiments revealed a functional bi-directional signaling between LHb neurons and astrocytes: stress first induces LHb neuronal activation, which triggers astrocytic calcium signals through LC- and α_1A_-AR-dependent noradrenergic signaling (Figure 3); in tandem, through gliotransmitters glutamate and ATP/Adenosine, LHb astrocytes prompt the second-phase of both local LHb neural activity and NE release (Figure 4 and 5). Through such a positive reinforcing loop, LHb-astrocytes facilitate stress-induced depressive-like behaviors (Figure 7).

**Figure 7.**
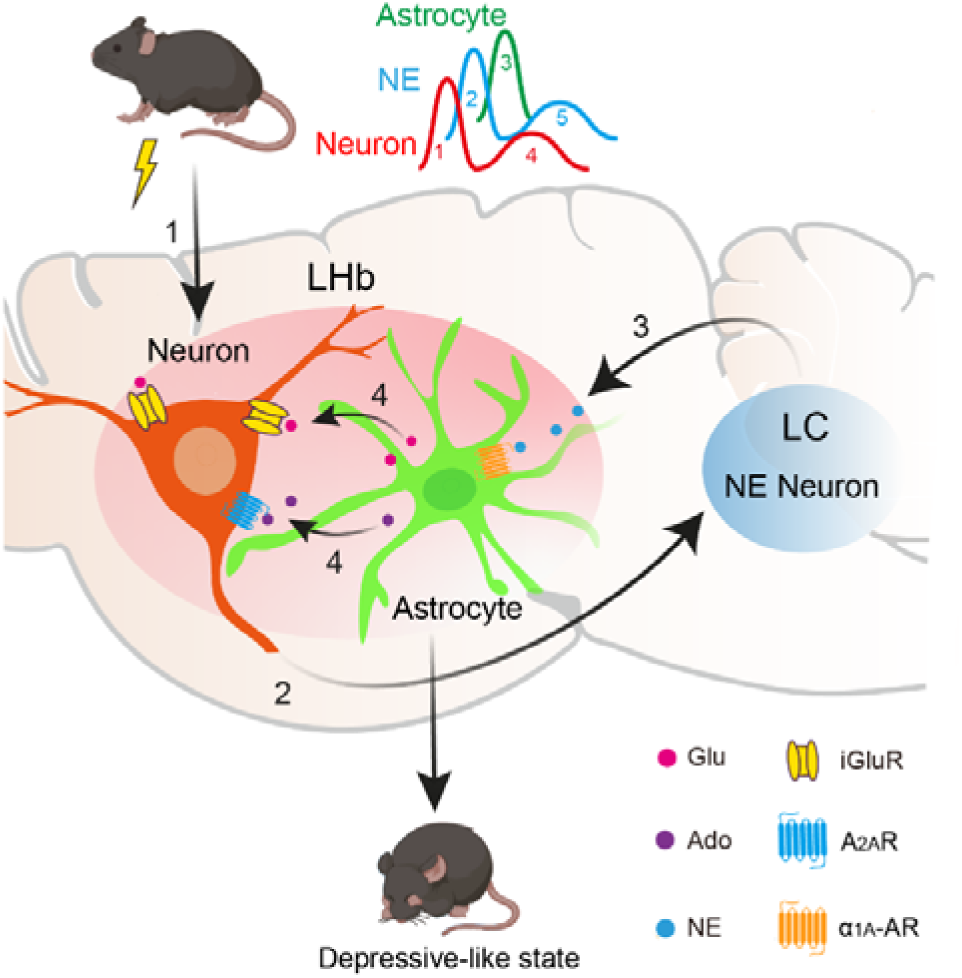
**LHb neuron-astrocyte synergy in mediating depression-like behaviors** (A) A model summarizing the LHb neuron-astrocyte synergy in mediating depression-like behaviors. Stress first activates LHb neurons, which in term recruit LC-NE neurons. NE release in LHb then triggers astrocytic calcium activities through α1AAR-dependent signaling. In tandem, through gliotransmitters glutamate (Glu) and ATP/Adenosine (Ado), LHb astrocytes prompt the second-phase of both local LHb neural activities and NE release. Through such a positive reinforcing loop and regulation of global gene expression, LHb astrocytes facilitate stress-induced depression-like behaviors.

### Functional implication and possible mechanisms underlying stress-induced biphasic calcium activities in LHb neurons

An important question in neurophysiology is how neurons maintain persistent activity in response to transient sensory stimuli. Previously several elegant mechanisms have been demonstrated, e.g. in the hypothalamus in maintaining a persistent internal emotional state^92^ or in the cortex in mediating working memory,^93,94^ involving slow-acting neuromodulators or recurrent neural network.^92–95^ Here our study reveals a new mechanistic foundation for a persistent neural dynamic pattern, in the context of FS-stress, driven by astrocyte signaling. In particular, we observed that LHb neuronal calcium signals exhibit a biphasic pattern, incorporating time-locked, fast first-phase transient and heavy-tailed, slow second-phase signals (Figure 2C and S3A-S3D). Importantly, the second-phase calcium signals, despite their smaller amplitude, lasted for many tens of seconds and had an AUC that was 3.46 ± 0.78 times that of the first phase (Figure 2C, 2D, S3D and S3E). Therefore, even more calcium enters into neurons during the second phase, which may activate calcium-dependent genes. These sets of genes are crucial for driving transcriptional programs, leading to persistent remodeling of neuronal network.^96,97^

In the other five brain regions examined, we did not observe such biphasic neural calcium activation in response to FS-stress (Figure S2A-S2E), suggesting that the above mechanism is region-specific. Such unique dynamics in the LHb may be accounted for by several potential mechanisms. First, the intrinsic properties of LHb neurons may be distinct. LHb neurons contain calcium-permeable AMPA receptors (CP-AMPARs)^15,17,98^ and pacemaker channels, such as hyperpolarization-activated cyclic nucleotide-gated (HCN) and T-type calcium channels.^18,99^ Through the glutamate-CP-AMPARs and adenosine-A2-receptors-coupled HCN channels,^86,100^ gliotransmitters released from LHb astrocytes may contribute to the second-phase neural calcium activation. Second, different types of gliotransmitters and different dynamics (e.g., onset and duration) of astrocytic activities in different brain regions may also contribute to this distinct neuronal response.

### Functional implication and possible mechanisms underlying stress-induced unique activities in LHb astrocytes

Compared with the five other brain regions recorded, astrocytic calcium dynamic in the LHb showed the earliest onset in response to FS-stress (Figure 1D-1G). This unique property may also be explained by several potential mechanisms. One possibility is that the strength of astrocytic coupling can potentially influence the speed of calcium propagation. Astrocytes form interconnected networks via gap junctions, enabling locally-coupled calcium activity among neighboring astrocytes.^35,47,101,102^ Heterogeneity of astrocytic gap-junction coupling in the central nervous system (CNS) may yield diverse calcium dynamics. Alternatively, different neuromodulator receptor signaling in astrocytes of different regions may account for varying time course and magnitude of calcium signals.^103^ For instance, in the context of FS-stress, astrocytes in auditory cortex are regulated by ACh,^67^ whereas in the LHb, they are regulated by NE (Figure 3H). Another possibility is that the sensitivity of astrocytes to NE may be differentially regulated by local neuronal activities. Supporting this, a previous study in visual cortex demonstrated that local neuronal activity serves as gain control to modulate the response of astrocytes to NE.^42^ On the other hand, our study demonstrates that local neural activity itself is not sufficient (Figure 3D-3F), and the long-range LHb-LC loop is required, to drive astrocytic calcium in the LHb (Figure 3G-3T).

The function of the stress induced LHb astrocytic activity may go beyond gene regulation in the local neuron-astrocyte network. Through the second-wave activation of LC-NE neurons, which have brain-wide projections,^104,105^ local LHb astrocytes may impact on global neural activity.

### Phasic and tonic NE release

Monoamine release is often subdivided into phasic (rapid kinetics) and tonic (slow kinetics) modes.^106–109^ Recent studies using biosensors have revealed both slowly ramping and phasic signals of monoamines in behaving mice, such as striatal dopamine release during reward-directed virtual reality and basal forebrain NE release under FS stress.^110,111^ During virtual reality experiments, tonic and phasic dopamine signals convey information of speed and changes in teleportation, respectively.^110^ These findings provide key insights into monoamine coding, suggesting that different modes of monoamine release may support different brain function.^106,112^

Here we found that NE release in the LHb during FS stress exhibited rapid phasic as well as slowly ramping tonic signals (Figure 3K-3L, S7A and S7C). The phasic NE release appears to be critical in activating LHb astrocytic calcium signals. Our recording showed that large increases in LHb astrocytic calcium activity were time-locked to the phasic NE signals (Figure 3M). Interestingly, although toward later shocks, the absolute amplitude of tonic NE signals was also high, as high as the amplitude of the early phasic signals, astrocytic calcium was still not induced (Figure S7). This suggests that to activate α1-adrenergic-receptor-dependent calcium signaling in LHb astrocytes, what matters is not only the amount but also the pattern of NE release. Different ligand-binding kinetics (e.g., the duration of dwell-time) can bias GPCR signaling toward the G-protein- biased or β-arrestin-biased pathways.^113,114^ In comparison to acute, sharp increase during phasic NE release, tonic stead-state increase of NE occurs over minutes time scale, which may induce bias towards β-arrestin-recruitment and receptor desensitization.^113^ It will be interesting to clarify the spatiotemporal organization and functional significance of these distinct NE signal patterns in the future.

As to the tonic NE signal, it displays the intriguing “ramp up” pattern over repeated footshocks (Figure 3K). We found that LHb astrocytes not only contribute to the second-phase phasic NE activity, but are also responsible for this ramp-up signal (Figure 5U and 5W). It will be very interesting to explore whether this increased basal NE activity may represent a change relevant for depression pathology, and through which mechanism astrocytes contribute to this change.

### Astrocyte function in behavioral regulation

Astrocytes were long considered passive bystanders of the brain. However, they are now emerging as important orchestrators of brain function and behaviors.^115,116^ Recent studies have highlighted the essential nature of a cohesive neuron-astrocyte program in influencing behavioral outcomes. Their coordinated activities span various processes including sensory information processing,^56,117^ brain metabolism,^118–121^ sleep regulation,^122^ memory consolidation,^123,124^ dominance behavior,^60^ and behavioral-state switching.^60,69,125–127^ These interactions are implicated in animal models of psychiatric disorders,^128,129^ including hyperactivity with attention deficit,^130^ obsessive-compulsive-like behavior,^55^ schizophrenia,^131^ drug addiction,^132^ and depression.^20,40,133^ Adding to this general theme, our results demonstrate that LHb astrocytes play an essential role in stress-induced depression-like behaviors. Furthermore, we reveal a dynamic interaction and information flow between neurons and astrocytes in this process. By dissecting the cellular and signaling pathways involved, we outline a stress-induced recurrent network defined by LHb neurons, LC-NE neurons and LHb astrocytes, highlighting LHb astrocytes as a critical signaling relay in this positive feedback loop. Through the broadcast by the LHb-LC neural circuit, local LHb astrocytes can then regulate global brain activity. Collectively, our findings provide a framework for understanding mechanisms of neuron-astrocyte interaction underlying stress-induced depression, with significant implications for the LHb-LC loop as a central hub in psychiatric disorders.

## Limitations of the study

One of the limitations is that the biphasic calcium activity of LHb neurons was derived from an analysis of bulk signals. Without recording at single-cell solution, we could not differentiate whether the two-phase activity involves the same or different subpopulations of LHb neurons. Miniscope/endoscope imaging in the LHb, which is technically highly challenging, should be explored in the future. Another limitation is that we did not provide experimental data to explain why LHb astrocytes respond most rapidly to stress. We have discussed multiple candidate mechanisms above, and further investigation is required to elucidate the underlying processes.

## Acknowledgments

We thank Yingzhuo Gou, Yang Xiong, Min Chen, Huafeng Zhang, Shiqi Wang and colleagues from Hu lab for assistance with experiments; Minmin Luo for providing the GfaABC_1_D-cOpn5-mCherry virus; Xiaowei Chen and Kuan Zhang for gifting the GfaABC_1_D-lck-GCaMP6f virus; Qin Han and Hangjun Wu at the Center of Cryo-Electron Microscopy for technical assistance with two-photon imaging; and Hui Li and Xiaoke Xie for advice on two-photon imaging. Some graphic components were created with BioRender. This work was supported by the STI2030-Major Projects (2021ZD0203000 and 2021ZD0203001), the National Natural Science Foundation of China (32130042, 31830032 and 82288101), the New Cornerstone Science Foundation to H.H., Key R&D Program of Zhejiang (2024SSYS0016), the Fundamental Research Funds for the Central Universities (2023ZFJH01-01 and 2024ZFJH01-01), the Non-profit Central Research Institute Fund of Chinese Academy of Medical Sciences (2023-PT310-01), and the Project for Hangzhou Medical Disciplines of Excellence, and Key Project for Hangzhou Medical Disciplines to H.H..

## Author Contributions

H.H., Q.X., and J.W. designed the study. Q.X., J.W., and, J.Z. conducted the behavioral pharmacology experiments and the related behavioral analysis. Q.X., and, J.W. performed the *in vivo* fiber photometry recording. J.W., and, Q.X. performed the *in vitro* two-photon imaging. Q.X. performed immunohistochemistry experiments with the assistance of J.Z., Y.T., X.J. Z.N. assisted J.W., and, Q.X. in analyzing two-photon imaging data. Y.L., J.F., and, Z.W. shared NE sensor and adenosine sensor. X.L., and, H.M. contributed to experimental design and discussions. H.H. supervised the project and wrote the manuscript with the assistance of Q.X. and J.W.

## STAR Methods

### KEY RESOURCES TABLE

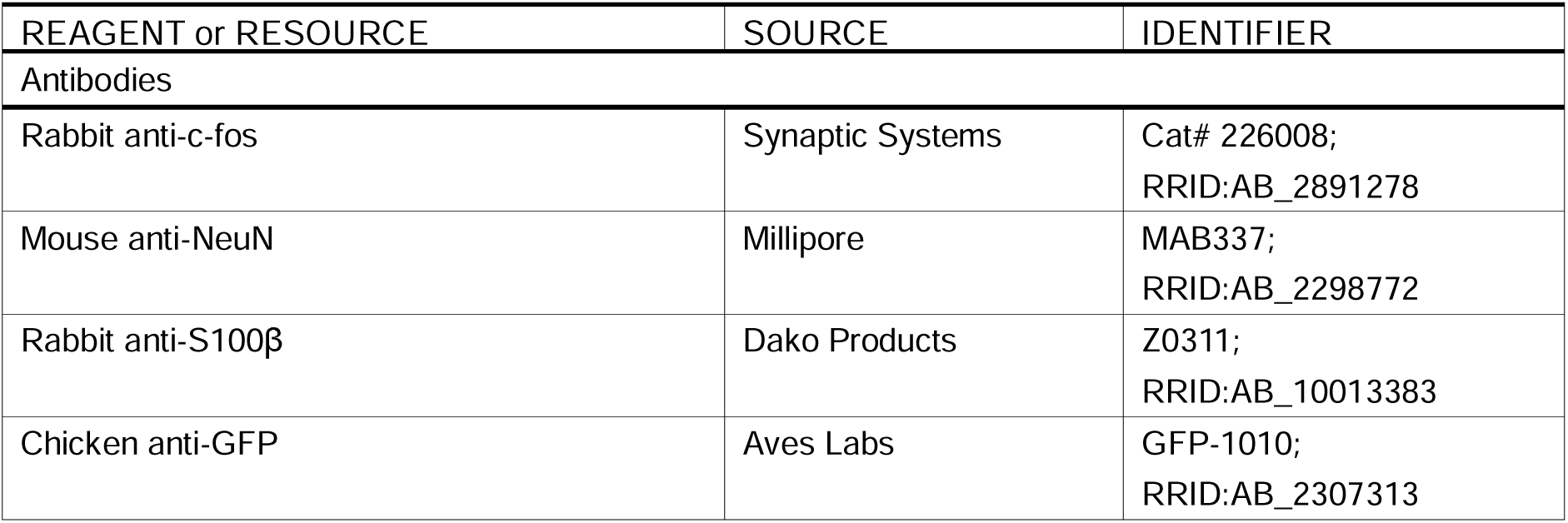
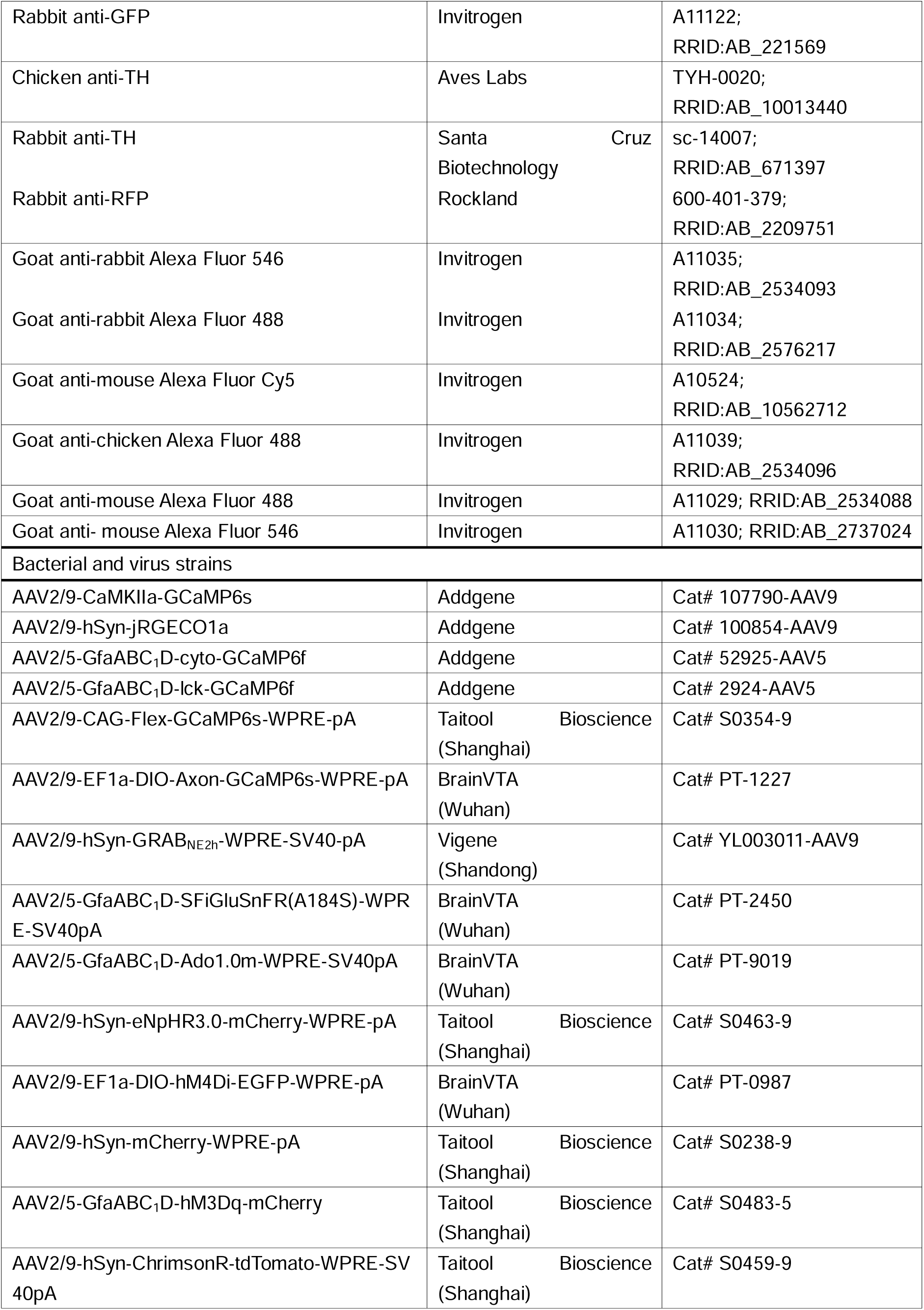
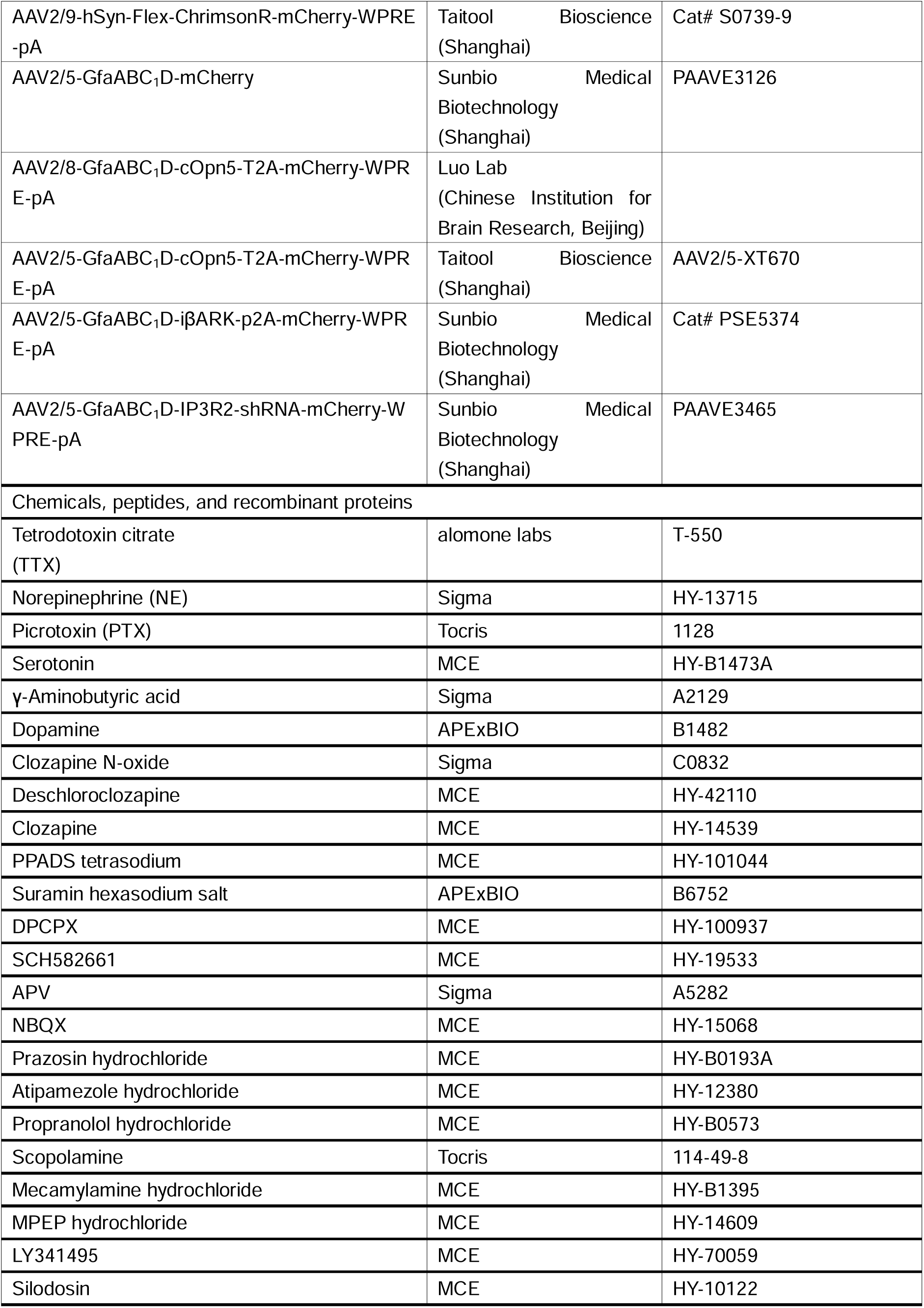
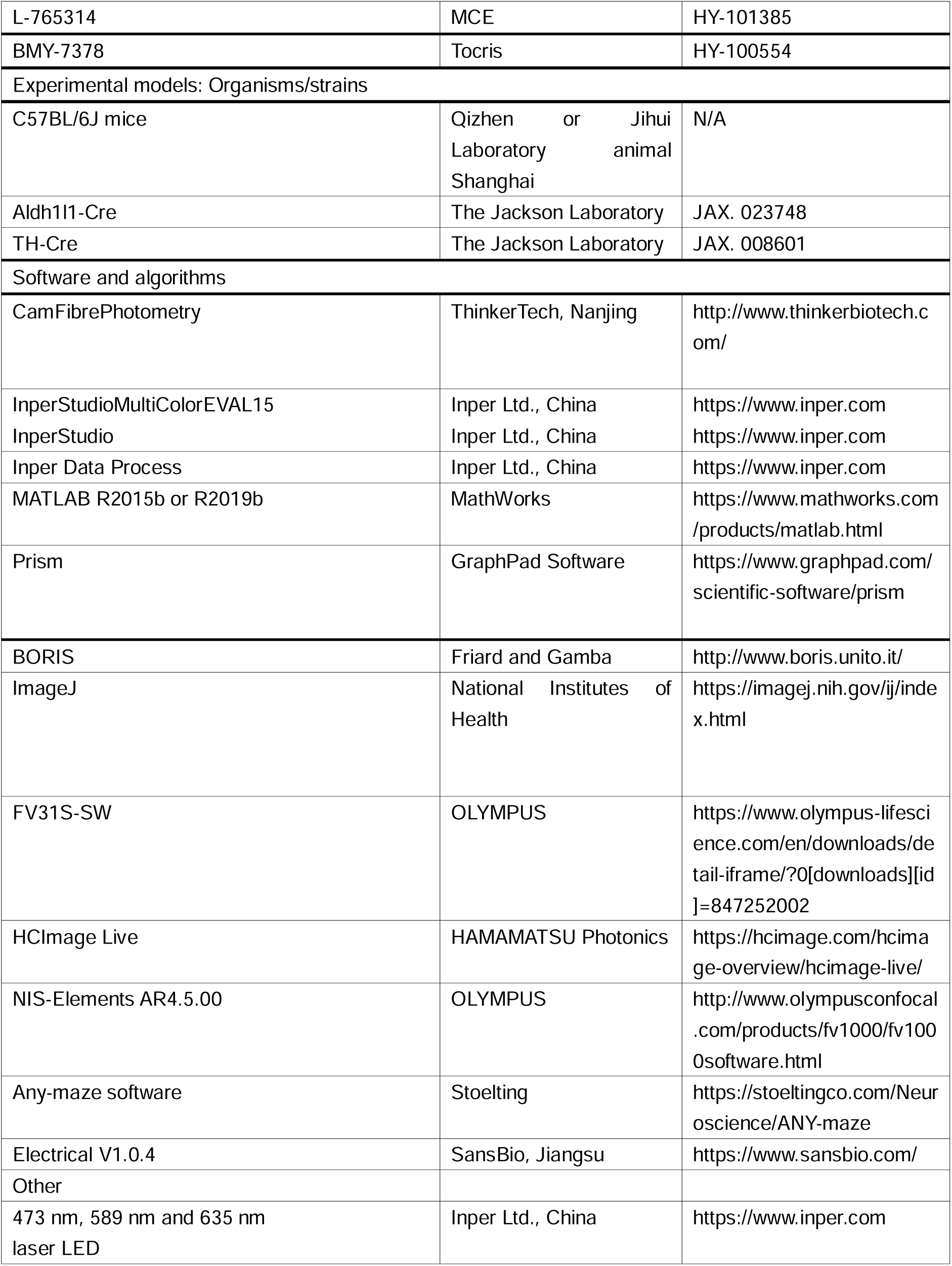
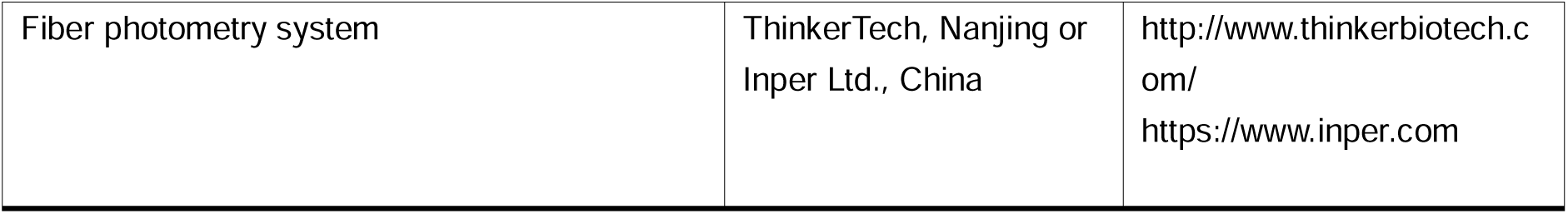

### EXPERIMENTAL MODEL AND SUBJECT DETAILS

#### Animals

Data for experiments were collected from male adult C57BL/6J strain mice (Qizhen or Jihui Laboratory animal), Aldh1l1-Cre (The Jackson Laboratory, JAX. 023748) and TH-Cre (The Jackson Laboratory, JAX. 008601) transgenetic mice, all over 8 weeks of age. Mice were housed in groups of four or five randomly under standard conditions (12-hour light/dark cycle) with food and water available *ad libitum*. Mice were habituated in the behavioral rooms for 0.5-1h before all the behavioral experimentations. All animal-related experimental procedures were under the guidelines of the Animal Care and Use Committee of the animal facility at Zhejiang University.

### METHOD DETAILS

#### Surgery and viral injection

Mice were deeply anaesthetized by 1% pentobarbital sodium (100mg/kg body weight) and head-fixed in a stereotactic frame (RWD Instruments). Virus was injected into the target brain regions (LHb: AP, - 1.72 mm from bregma; ML, ± 0.46 mm; DV, - 2.62 mm from the brain surface; mPFC: AP, + 2.43 mm from bregma; ML, ± 0.4 mm; DV, - 1.3 mm from the brain surface; LH: AP, - 0.90 mm from bregma; ML, ± 1.10 mm; DV, - 4.90 mm from the brain surface; STR: AP, + 0.80 mm from bregma; ML, ± 2.0 mm; DV, - 2.20 mm from the brain surface; BLA: AP, + 1.40 mm from bregma; ML, ±3.10 mm; DV, - 4.5 mm from the brain surface; HPC: AP, -2.20 mm from bregma; ML, ± 1.35 mm; DV, - 1.20 mm from the brain surface; LC: AP, -5.5 mm from bregma; ML, ± 0.9 mm; DV, - 3.40 mm from the brain surface) using a pulled glass pipette connected to a pressure microinjector (Picospritzer III, Parker). The injection needle remained in place for approximately 10 min before being withdrawn to prevent fluid reflux. For optic fiber implantation, a 200-μm optic fiber was placed 300 μm above the center of the viral injection site and cemented onto the skull with dental acrylic. For bilateral inhibition of the LHb neurons or multi-regions photometry recording, optic fiber was implanted to target brain regions (LHb: -1.72 mm AP, ±1.14 mm ML, -2.35 mm DV, and angled at 15 from the vertical in the lateral direction; LH: -0.9 mm AP, ±2.32 mm ML, -4.4 mm DV, and angled at 15 from the vertical in the lateral direction). After surgery, mice were allowed to recover from anesthesia on a heat pad.

After all experiments were completed, mice were transcardially perfused under deep anesthesia with 0.1 M phosphate buffered saline (PBS) followed 4% w/v paraformaldehyde (PFA) to verify the sites of virus injection and fiber placement. Brains were postfixed in 4% w/v PFA for 12 hours, followed by cryoprotection in a 30% w/v sucrose solution for 1∼2 days. The dehydrated brains were then sectioned into 40 μm thick coronal slices using a cryostat (Leica). The slices were counterstained with DAPI or Hoechst before imaging. Fluorescent image acquisition was performed using an Olympus Fluoview FV1000 confocal microscope. Only data from mice with correct virus injection site and optic location site were used.

For calcium recording experiments, the following viruses were used: AAV2/9-CaMKIIa-GCaMP6s (titre: 1.0 × 10^13^ vector genome (v. g.)/ml, dilution: 1: 8, 0.1μl into LHb, BLA, 0.2μl into LH, mPFC, STR, HPC, Addgene), AAV2/9-hSyn-jRGECO1a (titre: 1.0 × 10^13^ vector genome (v. g.)/ml, dilution: 1:3, 0.15μl into LHb, Addgene), AAV2/5-GfaABC_1_D-cyto-GCaMP6f (titre: 7.0 × 10^12^ vector genome (v. g.)/ml, 0.25μl into LHb, BLA, LH, mPFC, HPC, STR, Addgene), AAV2/5-GfaABC_1_D-lck-GCaMP6f (titre: 7.0 × 10^12^ vector genome (v. g.)/ml, 0.25μl into LHb, BLA, LH, mPFC, HPC, STR, Addgene), AAV2/9-CAG-Flex-GCaMP6s-WPRE-pA (titre: 1.66 × 10^13^ vector genome (v. g.)/ml, dilution: 1: 5, 0.2μl into LHb, Taitool Bioscience), AAV2/9-EF1a-DIO-Axon-GCaMP6s-WPRE-pA (titre: 5.14 × 10^12^ vector genome (v. g.)/ml, 0.2μl into LC, BrainVTA).

For sensor recording experiments, the following viruses were used: AAV2/9-hSyn-GRAB_NE2h_-WPRE-SV40-pA (titre: 1.0 × 10^13^ vector genome (v. g.)/ml, dilution: 1: 3, 0.2μl into LHb, Vigene), AAV2/5-GfaABC_1_D-SFiGluSnFR (A184S)-WPRE-SV40pA (titre: 1.0 × 10^13^ vector genome (v. g.)/ml, 0.2μl into LHb, BrainVTA), AAV2/5-GfaABC_1_D-ADO1.0m-WPRE-SV40pA (titre: 1.0 × 10^13^ vector genome (v. g.)/ml, 0.2μl into LHb, BrainVTA).

For manipulation experiments of neurons, the following viruses were used: AAV2/9-hSyn-ChrimsonR-tdTomato-WPRE-SV40pA (titre: 2.27 × 10^13^ vector genome (v. g.)/ml, dilution: 1: 10, 0.1 μl into LHb, Taitool Bioscience), AAV2/9-hSyn-eNpHR3.0-mCherry-WPRE-pA (titre: 1.76 × 10^13^ vector genome (v. g.)/ml, dilution: 1: 8, 0.1 μl into LHb, Taitool Bioscience), AAV2/9-hSyn-mCherry-WPRE-pA (titre: 2.49 × 10^13^ vector genome (v. g.)/ml, dilution: 1: 5, 0.1 μl into LHb, Taitool Bioscience), AAV2/9-hSyn-Flex-ChrimsonR-mCherry-WPRE-pA (titre: 1.30 × 10^13^ vector genome (v. g.)/ml, dilution: 1: 5, 0.15 μl into LC, Taitool Bioscience), AAV2/9-EF1a-DIO-hM4Di-EGFP-WPRE-pA (titre: 5.35 × 10^12^ vector genome (v. g.)/ml, 0.2μl per side bilateral into LC, BrainVTA).

For manipulation experiments of astrocytes, the following viruses were used: AAV2/5- GfaABC_1_D-hM3Dq-mCherry (titre: 1.45 × 10^13^ vector genome (v. g.)/ml, dilution: 1: 5, 0.2μl per side bilateral into LHb, plasmid from Addgene); AAV2/5- GfaABC_1_D-mCherry (titre: 2.08 × 10^13^ vector genome (v. g.)/ml, dilution: 1: 5, 0.2μl per side bilateral into LHb, Sunbio Medical Biotechnology); AAV2/8-GfaABC_1_D-cOpn5-T2A-mCherry-WPRE-pA (0.2μl per side bilateral into LHb, provided by M. Luo); AAV2/5-GfaABC_1_D-cOpn5-T2A-mCherry-WPRE-pA (0.2μl per side bilateral into LHb, Taitool Bioscience); AAV2/5-GfaABC_1_D-iβARK-p2A-mCherry-WPRE-pA (titre: 1.9 × 10^13^ vector genome (v. g.)/ml, 0.2μl per side bilateral into LHb, plasmid from Addgene); AAV2/5-GfaABC_1_D-IP3R2-shRNA-mCherry-WPRE-pA (titre: 2.61 × 10^13^ vector genome (v. g.)/ml, 0.2μl per side bilateral into LHb, Sunbio Medical Biotechnology).

#### In vivo fiber photometry recording

The calcium signals of multi-regions were simultaneously recorded using fiber photometry systems (ThinkerTech, Nanjing). A beam of 488 nm excitation light was delivered, and fluorescence signals were acquired at a sampling rate of 50 Hz. To minimize fluorescence signal bleaching, the laser intensity was adjusted to a low level (40 μW) at the tip of optic fiber.

The calcium signals of astrocytes and neurons were simultaneously recorded using dual-color fiber photometry recording systems (ThinkerTech, Nanjing; Inper Ltd., China). System delivered two excitation light sources, 470 nm and 580 nm, allowing simultaneous recording of red and green indicators at a frequency of 40 or 25 frames per channel per second. The light power was approximately 40 μW for the 470 nm wavelength and the 50 μW for the 580 nm wavelength at the tip of optic fiber.

During the recording of signals under the FS stress, the mice were placed into an FS chamber (SansBio, Jiangsu) with metal grid floor and subjected to unpredictable FS (1s, 0.4-1mA). A video camera was positioned above the chamber to track each mouse.

#### Optogenetic manipulations in fiber photometry recording

To synchronize optogenetic manipulation and photometry recording, we utilized fiber photometry recording systems combined with photostimulation (Inper Ltd., China). In the experiments involving the activation of LHb neurons, LHb-LC terminals and LC^TH^-LHb terminals, we applied 1s red-light pulses (635 nm, 2-5 mW at fiber tip, 5 ms or 10 ms pulse width). In the experiments involving inhibition of LHb neurons, we applied constant yellow light (589 nm, 10 mW at fiber tip). When the light was delivered, a trigger simultaneously sent a TTL pulse to the data recording systems, allowing verification of the exact time points when the light was turned on.

#### Analysis of fiber photometry data

Data were analyzed using the codes (e. g. OpSignal, from Thinker Tech Nanjing Biotech Co., Ltd. and Inper Ltd., China) based on MATLAB. The fluorescence responses were indicated by delta-F/F0 (calculated as (F-F0)/ F0). F0 represents the baseline average fluorescence signals in a 2-second-long period prior to the onset of FS stress or light on. Delta-F/F0 are presented as heatmaps and also as average plots with a shaded area indicating the SEM. For the quantification of rise and decay of signals, rise time was defined as latency from 10% peak signal timing to 80% peak signal timing. Decay time was defined as latency from peak timing decay to 50% peak activity timing. The peak of signals during response period (10s from the stimulus onset) was detected by finding a maximum response. For quantification of inflection point, average fluorescence signals evoked by FS stress for each mouse were used to define the point of decay slope change as the inflection point. For signal-phase analysis of neuronal calcium, phase1 was defined as the period from the onset of FS timing to the inflection point timing and phase2 was defined as the period from the inflection point timing to 10% peak activity timing. For astrocyte-loss-of-function experiments, the phases of neuronal calcium signals were defined as follows: phase1 was the period from the onset of FS to 2.5s, and phase2 was from 2.5 s to 40 s (It is difficult to identify the inflection point of neuronal calcium signals when inhibiting astrocytic calcium signaling). The quantification of AUC1 and AUC2 of NE signals evoked by FS excluded ramp values.

#### Drugs application in vivo recording

The following drugs were administered intraperitoneally (i.p.). Concentrations were as follows: Prazosin hydrochloride (4 mg/kg, MCE), Propranolol (40 mg/kg, MCE), Atipamezole hydrochloride (0.5 mg/kg, MCE), Scopolamine (1 mg/kg, Tocris), Mecamylamine hydrochloride (2.5 mg/kg, MCE), MPEP hydrochloride (20 mg/kg, MCE), LY341495 (2.5 mg/kg, MCE), Silodosin (5 mg/kg, MCE), L-765314 (5 mg/kg, MCE) and BMY-7378 (2.5 mg/kg, Tocris).

For inhibition of LC NE neurons, four weeks after injection of AAV2/9-EF1a-DIO-hM4Di-EGFP-WPRE-pA into LC, clozapine (1mg/kg, MCE) or saline was administered to TH-Cre mice by intraperitoneal injection.

For activation of LHb astrocytes, four weeks after injection of AAV2/5-GfaABC_1_D-hM3Dq-mCherry or AAV2/5-GfaABC_1_D-mCherry and AAV2/9-CaMKIIα-GCaMP6s into LHb, clozapine (0.25, 0.5, 1mg/kg, MCE) or saline was administered to mice by intraperitoneal injection.

For experiments of inhibitors and antagonists administration, each recording session was separated by 30-50 minutes to allow for drug effectiveness.

#### Acute brain slice preparation for imaging

Sagittal LHb slices were prepared from C57BL/6J or Aldh1l1-Cre mice with AAV virus injection for in vitro calcium imaging. Briefly, mice were deeply anesthetized with pentobarbital sodium and decapitated with sharp shears. The brains were sliced in ice-cold modified artificial CSF (ACSF) (oxygenated with 95% O_2_ and 5% CO_2_) containing the following (in mM): 220 sucrose, 2 KCl, 6 MgCl_2_, 0.2 CaCl_2_, 26 NaHCO_3_, 1.2 NaH_2_PO_4_, and 10 D-glucose. A vibratome (Leica2000) was used to cut 300 μm brain sections. The slices were allowed to equilibrate for 30 min at 34-36 °C in normal ACSF containing (in mM): 125 NaCl, 2.5 KCl, 2 CaCl_2_, 1 MgCl_2_, 25 NaHCO_3_, 1.25 NaH_2_PO_4_, and 25 D-glucose with 1 mM pyruvate added, continuously bubbled with 95% O_2_ and 5% CO_2_. Slices were allowed to recover for at least 1h in the same buffer until use. All slices were used within 5-8 hours of slicing.

#### In vitro imaging

Slice preparation was performed as described above. In neuronal calcium imaging and sensors imaging experiments, images were captured on an Olympus two-photon microscope (FVMPE-RS, Tokyo, Japan) equipped with a mode-locked Ti:Sapphire laser (MaiTai DeepSee, Spectra-Physics, San Francisco, USA) set at 920 nm and a water-immersion objective lens (25×, N.A. 1.05; Nikon, Tokyo, Japan). Regions with virus expression in the LHb were identified and full-frame images were acquired at 0.92 Hz. In astrocytic calcium imaging experiments, for optogenetic manipulation experiments, images were captured on an Olympus two-photon microscope (FVMPE-RS, Tokyo, Japan). For other experiments, images were captured on a digital CMOS camera (ORCA-Flash4.0 V3; HAMAMATSU and FL 9BW; TUCSEN) with a water-immersion objective lens (40×, N.A. 0.85). Full-frame images were acquired at 4 Hz. A constant flow of fresh buffer perfused the imaging chamber at all times.

#### Optogenetic manipulations in two-photon imaging

In optogenetic manipulation experiments, an optical fiber connecting a photostimulation system (Inper, China) was placed on the slices and delivered red light (635 nm, 5mW at fiber tip, 5ms or 10ms pulse width) for neuronal ChrimsonR activation, or blue light (473 nm, 60µW at fiber tip, 1s constant) for astrocytic cOpn5 activation.

#### Drug applications in vitro

The following drugs were applied in the bath: Norepinephrine (MCE, HY-13715), Phenylephrine (10 µM, Sigma), γ-Aminobutyric acid (300 µM, Sigma), Serotonin (20 µM, MCE), Dopamine (10 µM, APExBIO), Clozapine N-oxide (CNO, Sigma), Picrotoxin (100 µM; Tocris), PPADS tetrasodium (50 µM; MCE), Suramin hexasodium salt (75 µM ; APExBIO), DPCPX (300 nM; MCE), SCH582661 (100 nM; MCE), APV (50 µM; Sigma), NBQX (10 µM; MCE), Prazosin hydrochloride (10 µM; MCE), Atipamezole hydrochloride (1 µM; MCE), Propranolol hydrochloride (10 µM; MCE), Tetrodotoxin cltrate (TTX; 1 µM; alomone labs).

For experiments of inhibitors and antagonists bath, each slice imaging session was separated at least 10 min to permit drug penetration.

#### Analysis of imaging data

Calcium imaging analysis was conducted using FIJI (ImageJ) and MATLAB. Image XY drift was corrected using MATLAB. For CMOS camera imaging data (excluding Figure S6F, G), the entire field of view (FOV) was selected as the region of interest (ROI). In Figure S6F, G, the FOV was uniformly divided into 128x128 segments and the segments with calcium activities exceeding the threshold (delta-F/F0 > 10% in response to 5 µM NE) were selected as ROIs. For two-photon imaging data of sensors and astrocytic calcium signals, regions exhibiting fluorescence (including soma and process of astrocytes) during the experiment were selected as ROIs. For two-photon imaging data of neuronal calcium signals, the neuronal somatic regions were selected as ROIs. The fluorescence responses were indicated by delta-F/F0 (calculated as (F-F0)/ F0). F0 represents the baseline average fluorescence signals in a 10-second period prior to the onset of light on or a 60-second period prior to the drug application. Delta-F/F0 were presented as heatmaps and also as average plots with a shaded area indicating the SEM.

The recorded neurons show three modes of spontaneous calcium activity at resting conditions. We calculated the standard deviation (SD) of calcium fluorescence value and the SD of calcium-event intervals for each neuron during the resting conditions. Neurons were classified as silent if they showed no calcium events during recording (SD of fluorescence value < 30); Neurons were classified as regular if they showed periodic calcium events during recording (SD of fluorescence value > 30 and SD of calcium-event interval < 10); Neurons were classified as irregular if they showed irregular calcium events during recording (SD of fluorescence value > 30 and SD of calcium-event interval > 10).

For the recording of LHb neurons during astrocytic-hM3Dq activation, we calculated the mean fluorescence for each neuron during the control period (180-second period before CNO application) and during the CNO-responsive period (180-second period during CNO application). Neurons were identified as excited if their mean fluorescence during CNO-responsive period was at least 2.5-fold above during the control period. For the recording of LHb neurons during astrocyte-cOpn5 activation, photostimulations (473 nm, 60 µW at fiber tip, 1s constant) were performed 6-8 times. Neurons were identified as excited if they exhibited high response fidelity (> 80%). To quantify the effect of inhibitors and antagonists, neurons with stable responses (response fidelity > 80%, signal decay < 10%) to cOPN5 in normal ACSF were included in the analysis. We separately calculated the mean cOPN5-evoked calcium peak in normal ACSF (control) and in ACSF with inhibitors or antagonists added. Neurons were identified as inhibited if their mean calcium peak in the presence of inhibitors or antagonists decreased by at least 30% compared to the control.

#### Immunohistochemistry

In experiment of immunohistochemistry for c-Fos, four weeks after injection of AAV2/5-GfaABC_1_D-hM3Dq-mCherry or AAV2/5-GfaABC_1_D-mCherry into LHb, deschloroclozapine (0.5 mg/kg, MCE), clozapine (0.25 mg/kg, 0.5 mg/kg, 1 mg/kg, MCE) or saline was administered to mice by intraperitoneal injection. Two hours after administration, mice were sacrificed for immunohistochemistry of c-Fos. In experiment of immunohistochemistry for NeuN, S100β, GFP and RFP, mice were sacrificed, after all the experiments were completed.

Mice were deeply anesthetized with 1% pentobarbital sodium and perfused transcardially with 0.1 M phosphate buffer saline (PBS, pH = 7.4) followed by 4% paraformaldehyde in PBS. Brains were removed and postfixed overnight and dehydrated in 30% sucrose in PBS. Coronal brain sections (40 μm) were serially cut and divided for 6 interleaved sets.

The antibodies used were rabbit anti-c-Fos (1:2000, SYSY), mouse anti-NeuN (1:500, Millipore), rabbit anti-S100β (1:1000, Dako Products), chicken anti-GFP (1:2000, Aves Labs), rabbit anti-GFP (1:2000, Invitrogen), chicken anti-TH (1:1000, Aves Labs), rabbit anti-TH (1:1000, Santa Cruz Biotechnology), rabbit anti-RFP (1:1000, Rockland); Alexa Fluor 546 goat anti-rabbit IgG, Alexa Fluor 488 goat anti-rabbit IgG, Alexa Fluor Cy5 goat anti-mouse IgG, Alexa Fluor 488 goat anti-chicken IgG, Alexa Fluor 488 goat anti-mouse IgG, Alexa Fluor 546 goat anti-mouse IgG (all 1:1,000, Invitrogen).

Slices for checking the injection site were counterstained with Hoechst in the final incubation step. Fluorescent image acquisition was performed with an Olympus Fluoview FV1000 confocal microscope.

#### Behavioral assays

##### Foot-shock stress (FS)

In FS-induced depression-like behaviors protocol, FS was performed as previously described.^16,91^ Mice were placed into a standard FS chamber (SansBio, Jiangsu) with metal grid floor and habituated to the new environment for 5 min. During a 20-min FS session, mice were subjected to 20 unpredictable FS (1 mA, 1s) with an intershock interval of 45-75s. In subthreshold FS protocol, mice were subjected to 6 unpredictable FS (1 mA, 1s) with an intershock interval of 45-75s. For activation of LHb astrocytic Gq pathway, Deschloroclozapine (0.5 mg/kg, MCE) was administered intraperitoneally (i.p.) 10 min before FS.

##### Forced swim test (FST)

FST was performed as previously described.^18,134^ Mice were gently and individually placed in a cylinder (12 cm diameter, 25 cm height) of water (23-24 °C) and swam for 6 min under normal light (100-200 lux). Water depth (15 cm) was set to prevent mice from touching the bottom with their tails or hind limbs. The entire test lasted for 6 min. A camera was set at the side of the cylinder to record the behaviors. The immobile duration during the last 4-min test were counted offline by an experienced observer blinded to the animal treatments. Immobility was defined by animals remaining floating or motionless with only small and necessary movements for keeping balance.

##### Sucrose preference test (SPT)

SPT was conducted as previously described.^18,134^ Animals were single housed and habituated to two bottles of drinking water for 2 days, followed by two bottles of 2% sucrose for 2 days. After habituation, the preference for any specific bottle was checked. Only mice without basal preference (between 25-75%) were used. Mice with basal preference of one bottle on the last day of habituation (below 25% or above 75%) were excluded. Mice were then water deprived for 24 hours. In the test phase, mice exposed to one bottle of 2% sucrose and one bottle of water for 2 hours in the dark phase. The positions of 2 bottles were switched after 1hour. The sucrose preference was calculated as the average of consumption of sucrose/ total consumption of water and sucrose during the 2 hours.

### Statistical analysis

All data are shown as mean ± SEM. Statistical analyses were done with Prism 6 (GraphPad) or MATLAB. By pre-established criteria, values were excluded from analyses if virus expression was poor or optic fiber location was out of the interested region. The data were analyzed by Student’s test for Gaussian distributions, while Mann-Whitney test for non-Gaussian distributions. Results were considered statistically significant when the P value < 0.05. More details are provided in the Table S1.

## Supplementary figure legends

**Figure S1.**
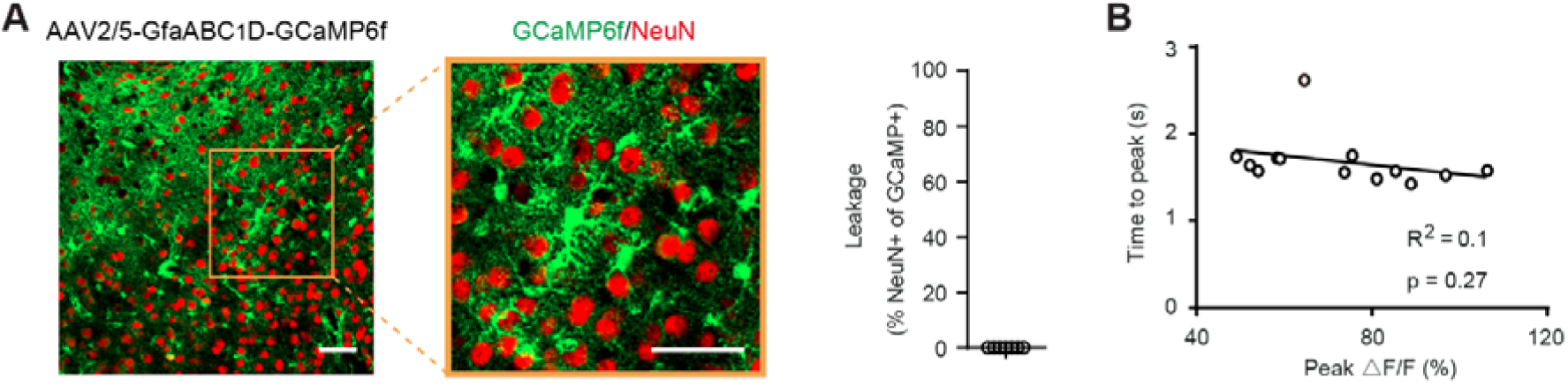
Foot-shock-stress Induces Most Rapid Calcium Response in LHb Astrocytes, Related to Figure 1 (A) GfaABC_1_D-promoter-driven GCaMP is not expressed in neurons. Left and middle: representative images of LHb brain slices stained with antibodies against GFP (indicating GCaMP-positive cells, green) and NeuN (neuronal marker, red). Scale bar, 50 μm. Right: bar graph showing percentage of GCaMP-positive cells that are NeuN-positive (n = 7 slices from 4 mice). (B) Scatterplots showing the relationship between peak amplitude and time to peak of LHb astrocytic calcium signals evoked by FS. Each circle represents one mouse. *P < 0.05; **P < 0.01; ***P < 0.001; ****P < 0.0001; n.s., not significant. Data are represented as mean ± SEM.

**Figure S2.**
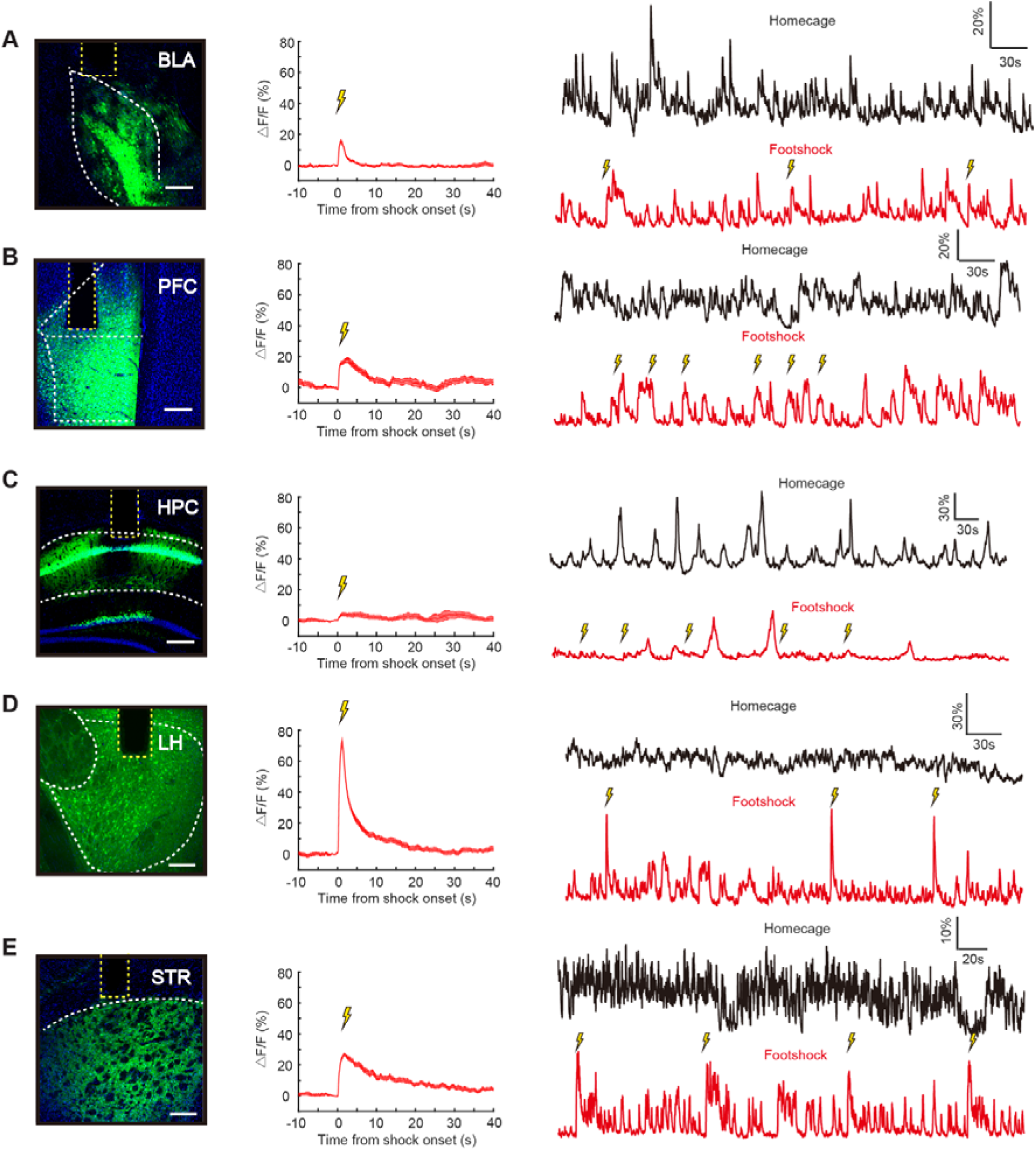
Neuronal Calcium Response Evoked by Foot-shock-stress in Other Five Brain Regions, Related to Figure 2 (A-E) Left: Illustration of viral expression and optic fiber placement (indicated by the yellow dotted line) in BLA (A), mPFC (B), HPC (C), LH (D) and STR (E), the boundary of which is outlined by the white dotted line. Green, GCaMP6s; blue, Hoechst. Scale bars, 100 μm. Middle: Average delta F/F ratio of neuronal calcium signals from BLA (A, n = 3 mice), mPFC (B, n = 7 mice), HPC (C, n = 7 mice), LH (D, n = 3 mice) and STR (E, n = 3 mice) aligned to the onset of FS. Solid lines indicate mean and shaded areas indicate SEM. Right: representative traces of recorded neuronal calcium signals from the 5 CNS regions during homecage (top) and continuous FSs (bottom).

**Figure S3.**
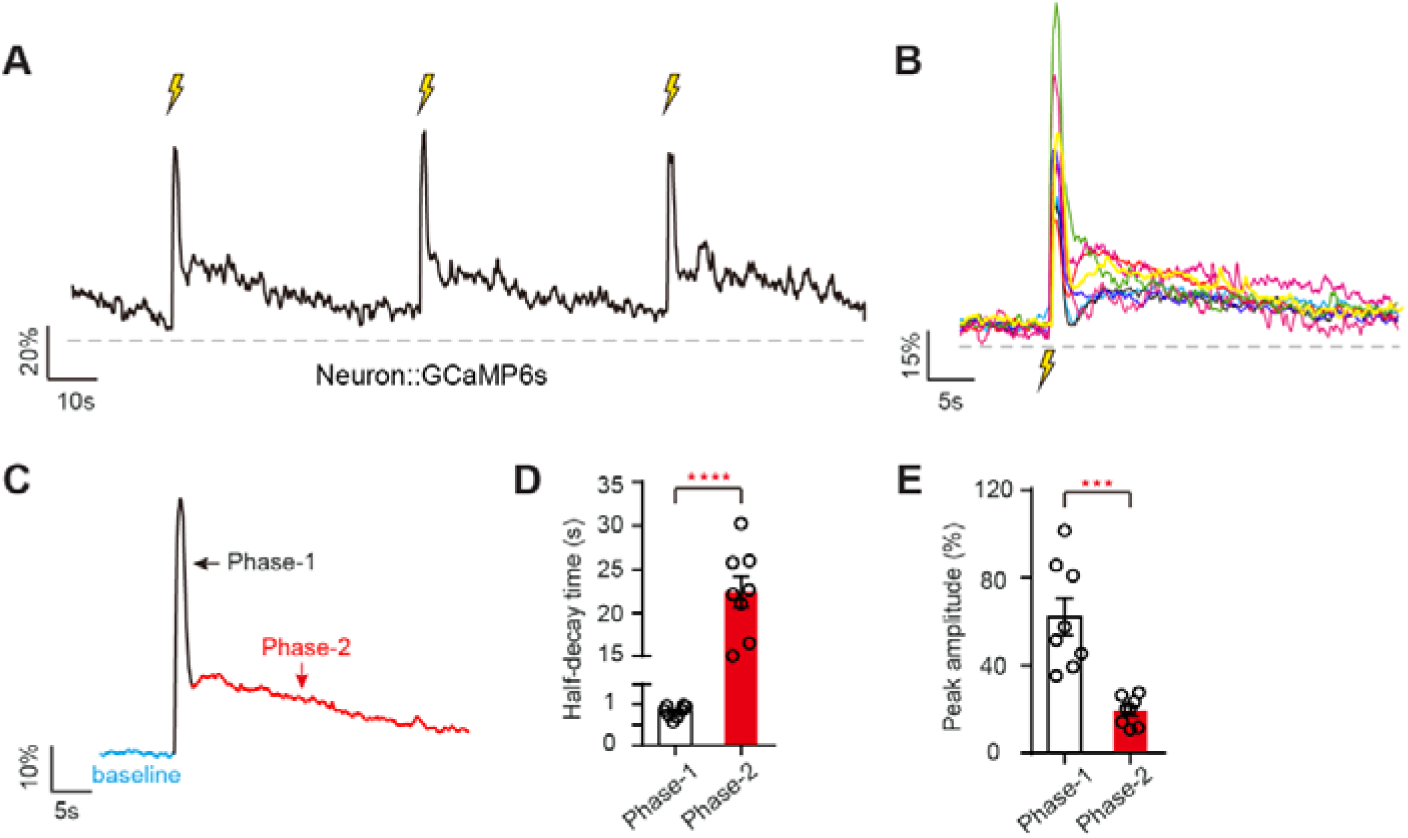
Foot-shock-stress Induces Biphasic Calcium Activities in LHb Neurons, Related to Figure 2 (A) Representative traces of recorded neuronal calcium signals in LHb over three continuous FSs. Scale bars, 20% delta F/F, 10s. (B) Alignment of FS-onset plots of average calcium signals of recorded neurons from individual animal during FS (n = 8 mice). Each circle represents one mouse. (C) Illustration of the segmented phases of FS-evoked LHb neuronal calcium signals. (D) Average half-decay time in phase-1 and phase-2 of FS-evoked neuronal calcium signals. Each circle represents one mouse. (E) Average peak amplitude in phase-1 and phase-2 of FS-evoked neuronal calcium signals. Each circle represents one mouse. *P < 0.05; **P < 0.01; ***P < 0.001; ****P < 0.0001; n.s., not significant. Data are represented as mean ± SEM.

**Figure S4.**
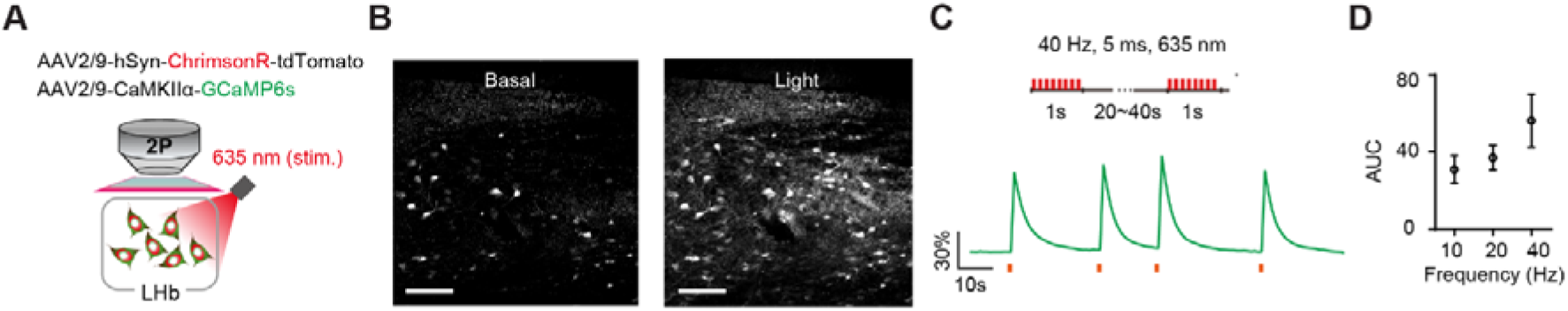
Optogenetic Activation of ChrimsonR-expressing Neurons Induces Neuronal Calcium in LHb Slice, Related to Figure 3 (A) Schematic illustrating optogenetic activation of ChrimsonR-expressing neurons while recording two-photon images of LHb neuronal calcium responses. (B) Calcium fluorescence images of LHb neurons before (B, left) and after (B, right) optogenetic activation of LHb neurons (1s, 40 Hz). (C) Neuronal calcium responses to optogenetic activation of LHb neurons (1s, 40 Hz) in LHb slices. (D) Mean neuronal calcium responses to varying stimulation frequencies in LHb slices.

**Figure S5.**
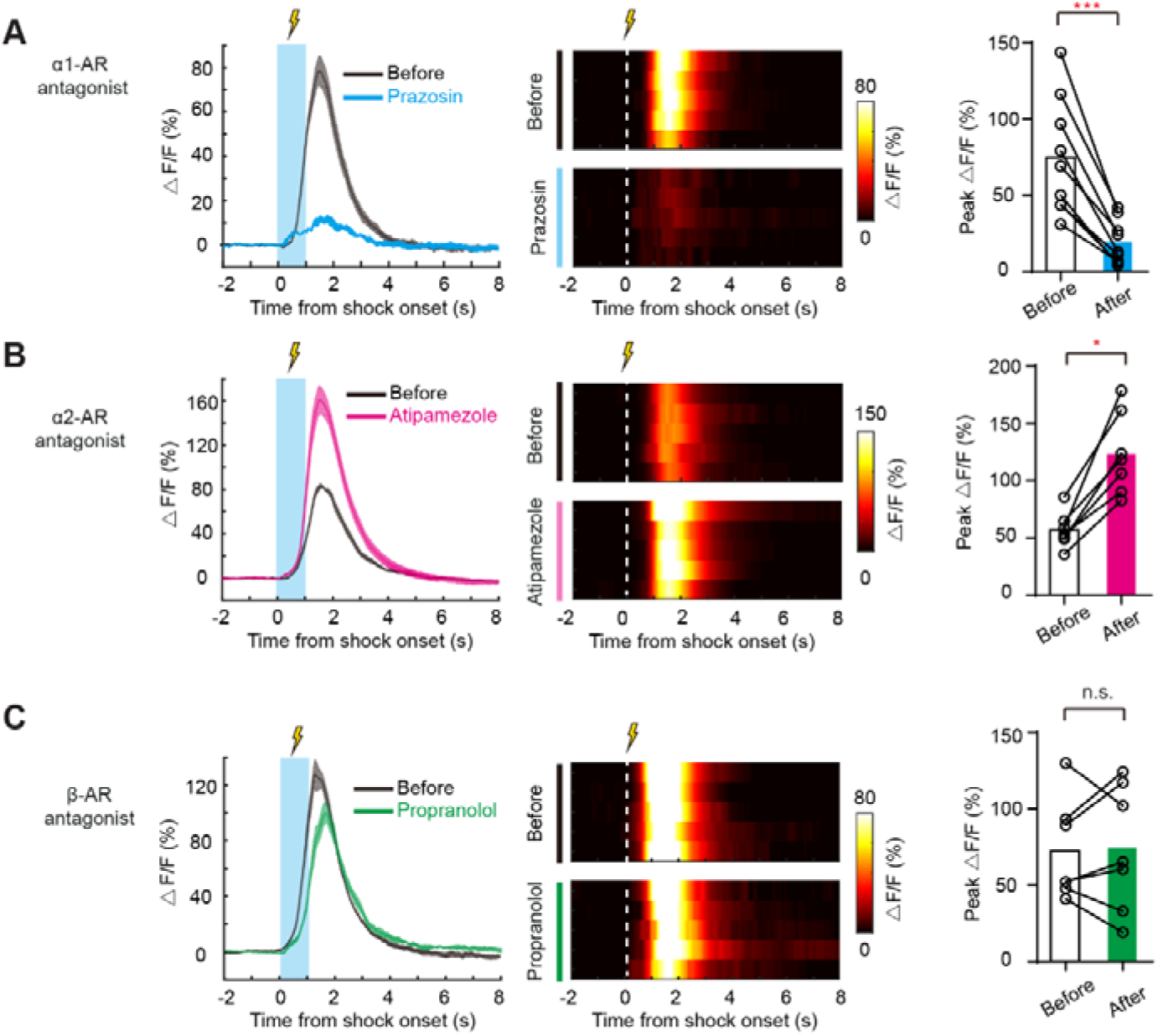
Calcium Activities Evoked by FS in LHb Astrocytes Require Noradrenergic Signaling, Related to Figure 3 (A-C) Plots of averaged delta F/F ratio (left) and heatmap (middle) of FS-evoked calcium signals in the LHb astrocytes before and after i.p. injection of Prazosin (α1-AR antagonist, A), Atipamezole (α2-AR antagonist, B) and Propranolol (β-AR antagonist, C) aligned to the FS onset. Bar graph showing effects of NE receptor antagonists on FS-evoked calcium signals in LHb astrocytes (right). Each circle represents one mouse. *P < 0.05; **P < 0.01; ***P < 0.001; ****P < 0.0001; n.s., not significant. Data are represented as mean ± SEM.

**Figure S6.**
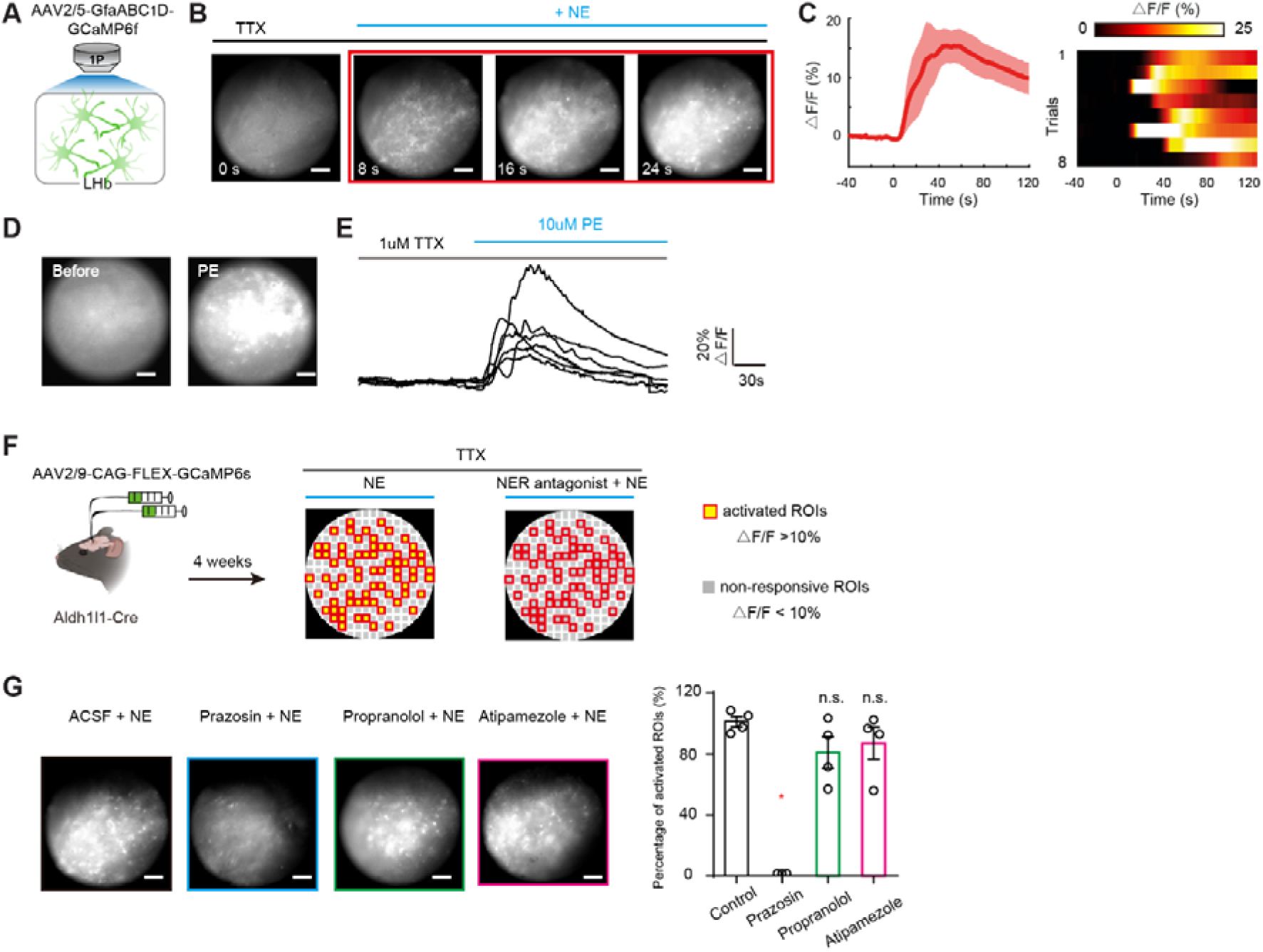
NE Triggers Calcium Signals in LHb Astrocytes via **α**1-AR, Related to Figure 3 (A) Schematic illustrating in vitro calcium imaging of LHb astrocytes in slices. (B) Calcium fluorescence images of LHb astrocytes showing calcium increase at different time points after NE application (n = 8 slices from 8 mice). Scale bar: 50 μm. (Experiments were performed in TTX). (C) Plots of delta F/F ratio (left) and heatmap (right) of calcium signals in LHb astrocytes induced by NE (n = 8 slices from 8 mice). Solid lines indicate mean and shaded areas indicate SEM. (D, E) Calcium fluorescence images of LHb astrocytes before (D, left) and after (D, right) application of phenylephrine (PE, α1-AR agonist). Traces (E) of LHb astrocytic calcium signals before and after PE application (n = 6 slices from 4 mice). Scale bars, 20% delta F/F, 30s. (Experiments were performed in TTX). (F) Schematic illustrating viral injection of Cre-dependent GCaMP6s in LHb in Aldh1l1-Cre mice and in vitro calcium imaging of LHb astrocytes in response to NE after application of NE receptor antagonist. ROI detection using a grid array (see STAR Methods for details). (G) Representative calcium fluorescence images (left) and bar graph (right) showing effects of Prazosin (α1-AR antagonist), Propranolol (β-AR antagonist) and Atipamezole (α2-AR antagonist) on LHb astrocytic calcium signals evoked by NE. Scale bar: 50 μm. Each circle represents one slice. *P < 0.05; **P < 0.01; ***P < 0.001; ****P < 0.0001; n.s., not significant. Data are represented as mean ± SEM.

**Figure S7.**
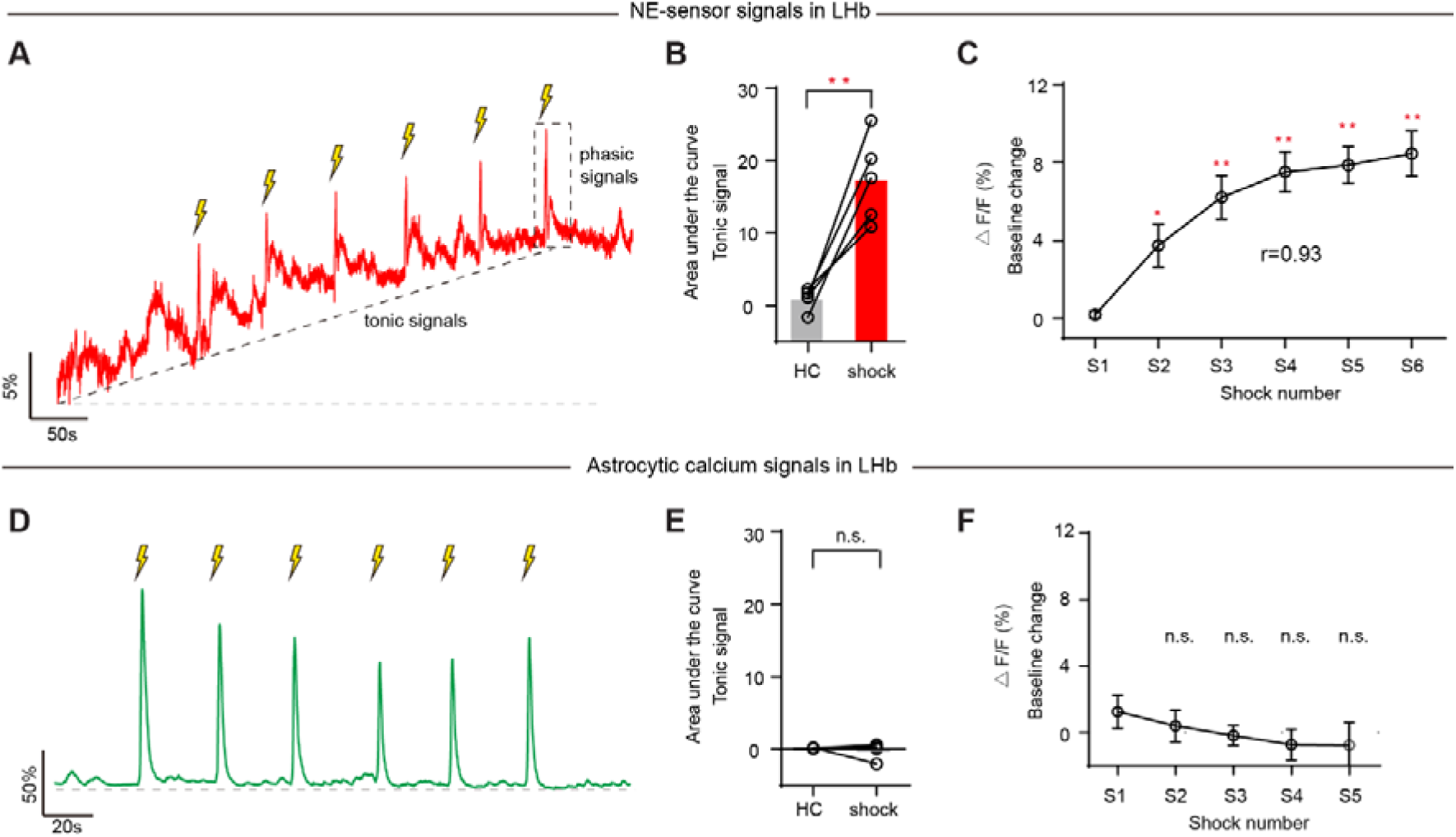
LHb NE Sensor and Astrocytic Calcium Signals Induced by FS, Related to Figure 3 (A) Example trace of LHb-NE-sensor signals over multiple FSs. Scale bars, 5% delta F/F, 50s. (B) AUC of basal (tonic) signal of LHb NE sensor in homecage (grey) and in response to FS (red). Each circle represents one mouse. (C) Plots showing basal (tonic) signal change of LHb NE sensor in response to FS. (D) Example trace of LHb astrocytic calcium signals over multiple FSs. Scale bars, 5% delta F/F, 50s. (E) AUC of basal signal of LHb astrocytic calcium in homecage (grey) and in response to FS (red). Each circle represents one mouse. (F) Plots showing basal signal change of LHb astrocytic calcium in response to FS. *P < 0.05; **P < 0.01; ***P < 0.001; ****P < 0.0001; n.s., not significant. Data are represented as mean ± SEM.

**Figure S8.**
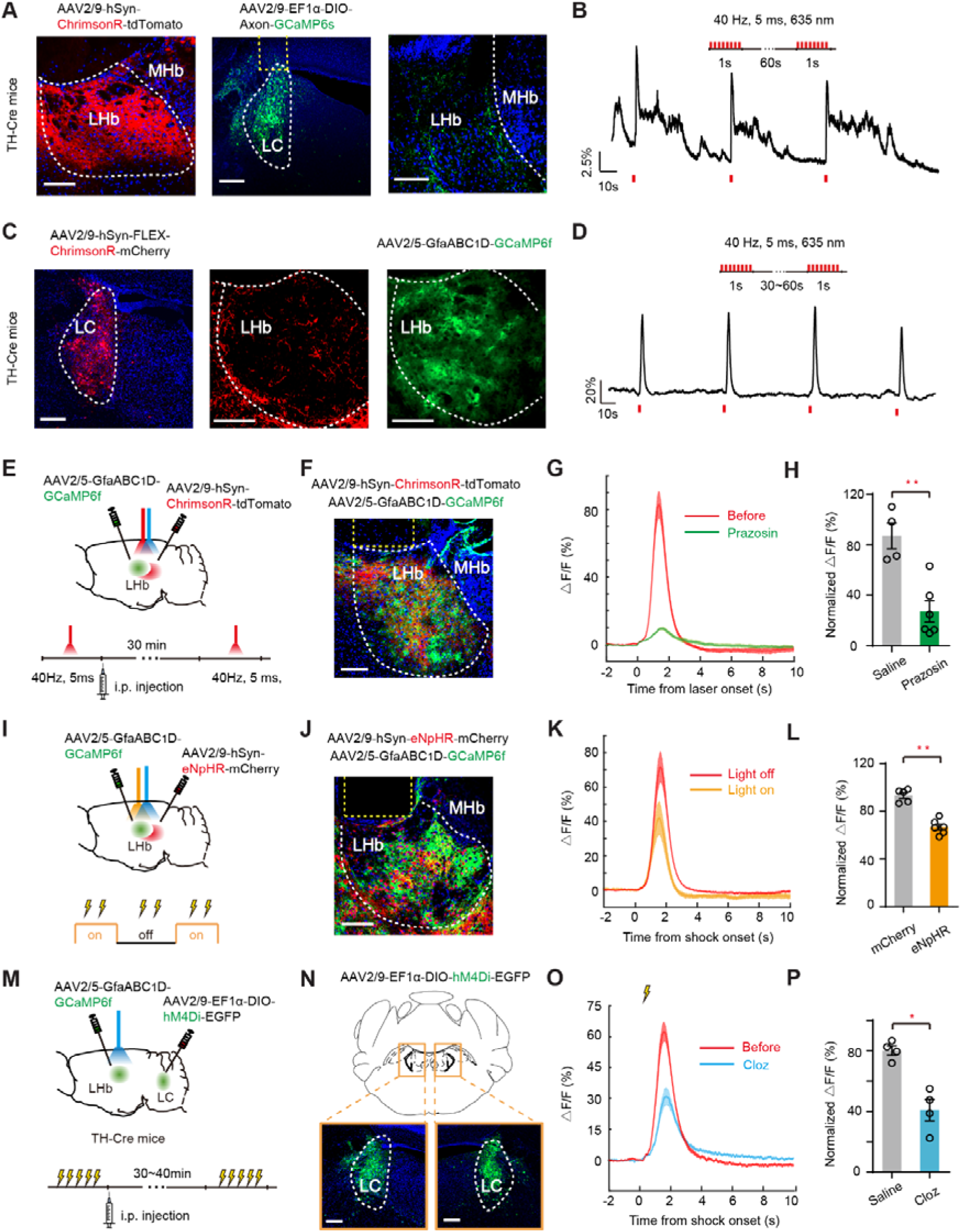
Effects of manipulating LHb and LC-NE Neurons on LHb Astrocytic Calcium, Related to Figure 3 (A, B) Effects of activating LHb-LC terminals on LC-NE neurons. (A) Representative images showing viral expression of ChrimsonR in LHb neurons (left), GCaMP6s in LC-NE neurons (middle) and LC-NE neuron terminals in LHb (right, stained with antibody against GFP) in TH-Cre mice. Scale bar, 100 μm. (B) Calcium responses of LC-NE neurons to optogenetic activation of LHb-LC terminals (1s, 40 Hz) in vivo. (C, D) Effects of activating LC^TH^-LHb terminals on LHb astrocytes. (C) Representative images showing viral expression of ChrimsonR in LC NE-neurons (left), LC^TH^-LHb terminals (middle, stained with antibody against RFP) and GCaMP6f in LHb astrocytes (right) in TH-Cre mice. Scale bar, 100 μm. (D) Calcium responses of LHb astrocytes to optogenetic activation of LC^TH^-LHb terminals (1s, 40 Hz). (E-H) Effects of Prazosin (α1-AR antagonist) on LHb astrocytic calcium signals evoked by LHb neuron activation in vivo. (E) Schematic illustrating optogenetic activation of LHb neruons and fiber photometry recording of LHb astrocytes. (F) Representative image showing viral expression of ChrimsonR in LHb neurons and GCaMP6f in LHb astrocytes. Scale bar, 100 μm. (G) Plots of averaged delta F/F ratio of LHb astrocytic calcium signals induced by LHb neurons before and after i.p. injection of Prazosin (α1-AR antagonist) aligned to the laser onset. Solid lines indicate mean and shaded areas indicate SEM. (H) Bar graph showing the effects of Prazosin (α1-AR antagonist) on astrocytic calcium signals evoked by LHb neurons. Each circle represents one mouse. (I-L) Effects of inhibiting LHb neurons on FS-evoked calcium signals in LHb astrocytes. (I) schematic illustrating optogenetic inhibition of LHb neruon and fiber photometry recording of LHb astrocytes in response to FS. (J) Representative image showing viral expression of eNpHR in LHb neurons and GCaMP6f in LHb astrocytes. Scale bar, 100 μm. (O) Plots of averaged delta F/F ratio of FS-evoked calcium signals in LHb astrocytes during light off and light on aligned to FS onset. Solid lines indicate mean and shaded areas indicate SEM. (L) Bar graph showing effects of inhibiting LHb neurons on FS-evoked calcium signals in LHb astrocytes. Each circle represents one mouse. (M-P) Effects of inhibiting LC-NE neurons on FS-evoked calcium signals in LHb astrocytes. (M) Schematic illustrating chemogenetic inhibition of LC-NE neurons and fiber photometry recording of LHb astrocytes in response to FS. (N) Representative image showing viral expression of hM4Di in LC-NE neurons. Scale bar, 100 μm. (K) Plots of averaged delta F/F ratio of FS-evoked calcium signals in LHb astrocytes before and after i.p. injection of clozapine (Cloz) aligned to FS onset. Solid lines indicate mean and shaded areas indicate SEM. (P) Bar graph showing effects of inhibiting LC NE-neurons on FS-evoked calcium signals in LHb astrocytes. Each circle represents one mouse. *P < 0.05; **P < 0.01; ***P < 0.001; ****P < 0.0001; n.s., not significant. Data are represented as mean ± SEM.

**Figure S9.**
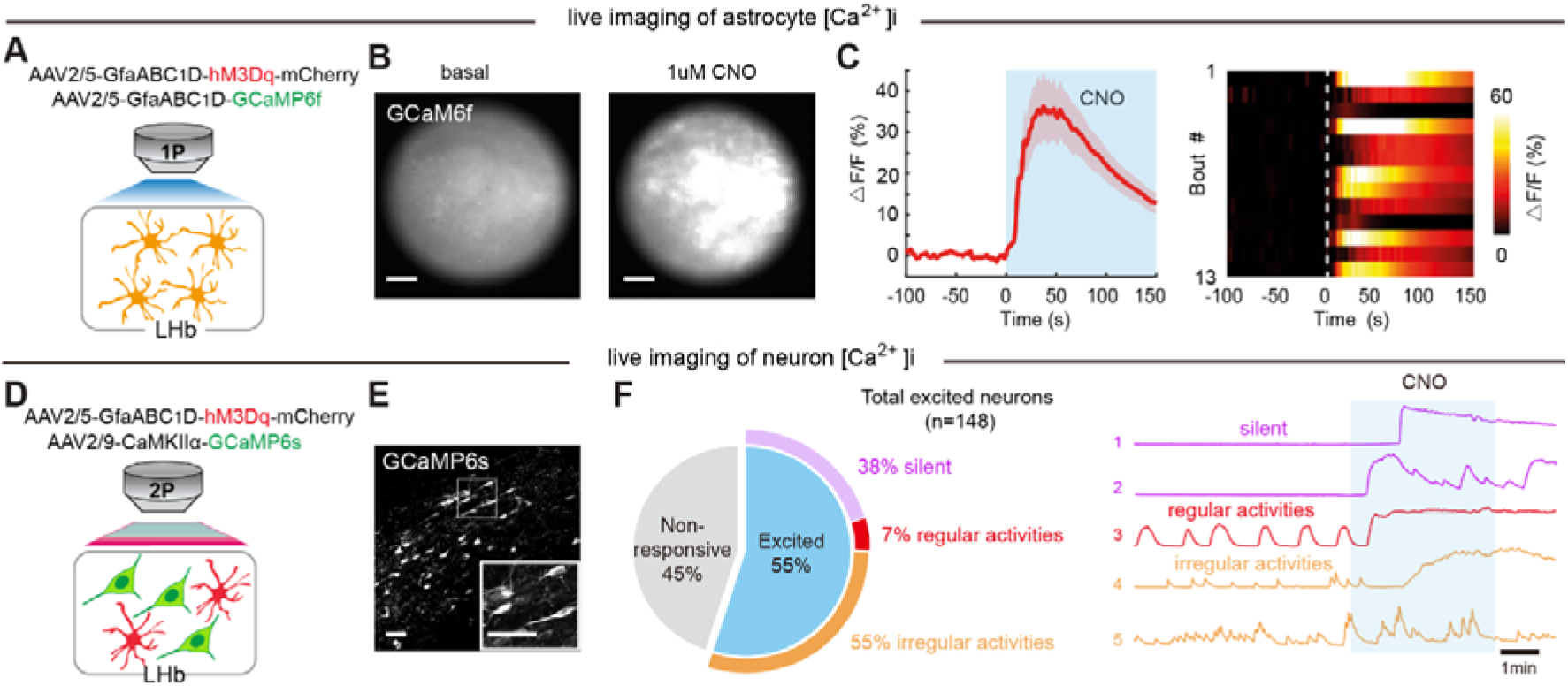
Chemogenetic Activation of Astrocytic Gq Pathway in LHb Slice, Related to Figure 4 (A, B) Schematic (A) and representative calcium fluorescence images (B) illustrating chemogenetic activation of hM3Dq-expressing astrocytes while recording calcium fluorescence images of LHb astrocytic calcium responses. Scale bar: 50 μm. (C) Plots of delta F/F ratio (left) and heatmap (right) of calcium signals in LHb astrocytes before and after CNO application (n = 3 mice). Solid lines indicate mean and shaded areas indicate SEM. (D, E) Schematic (D) and representative two-photon image (E) illustrating chemogenetic activation of hM3Dq-expressing astrocytes while recording two-photon image of LHb neuronal calcium responses. Scale bar: 50 μm. (F) Pie chart illustrating percent abundance of excited neurons by astrocytic hM3Dq activation (left, n = 3 slices from 2 mice). Colored outer circle indicates percentage of three types (according to baseline activity) among astrocytic-hM3Dq-excited neurons. Representative raw calcium traces from the three types of astrocytic hM3Dq-excited neurons (right).

**Figure S10.**
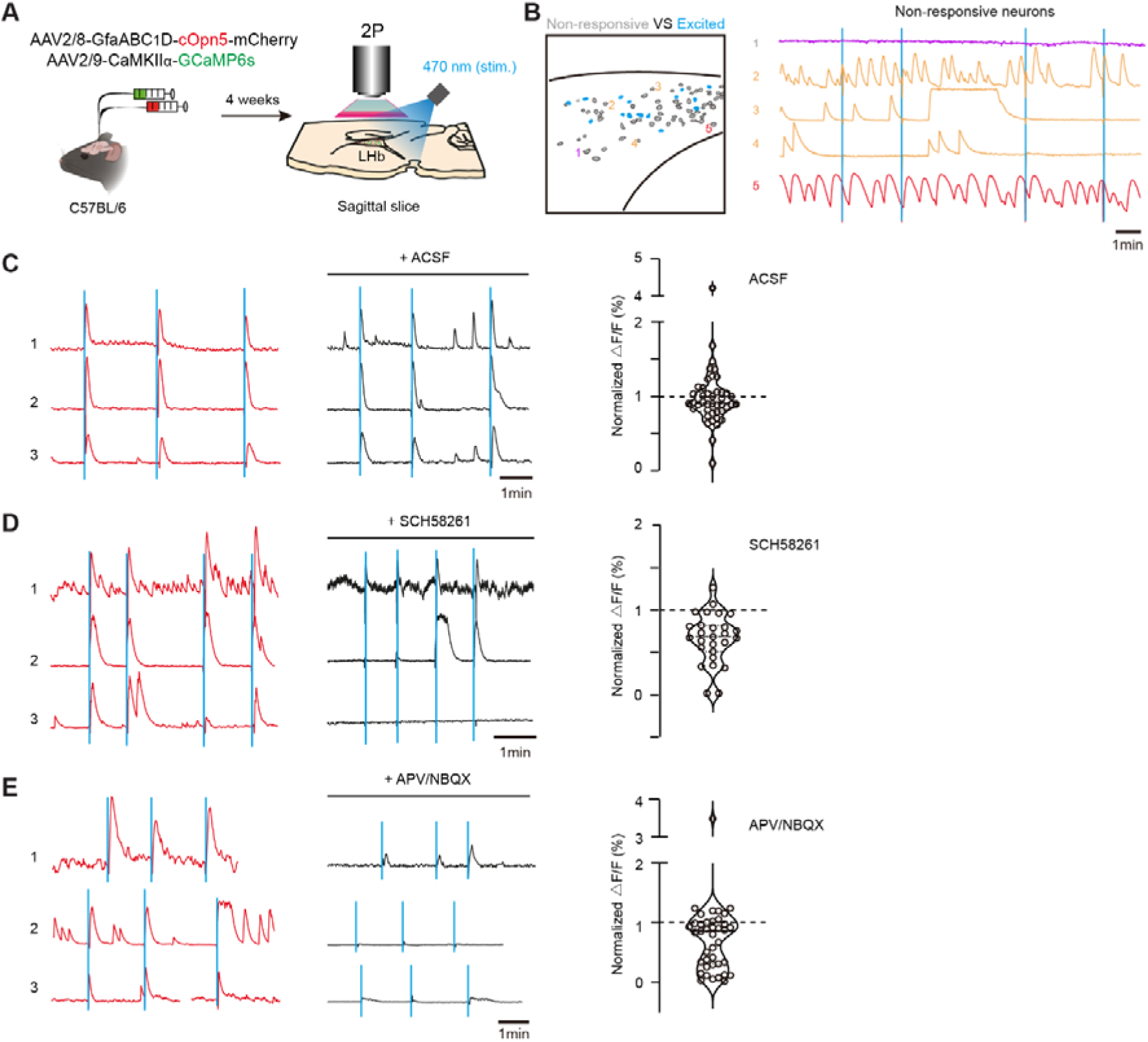
Optogenetic Activation of Astrocytic Gq Pathway in LHb Slice, Related to Figure 4 (A) Schematic illustrating optogenetic activation of cOpn5-expressing astrocytes while recording two-photon images of LHb neuronal calcium responses. (B) Left: example spatial map of astrocytic cOpn5-excited and non-responsive neurons in the LHb. Right: representative raw calcium traces from the three types of non-responsive neurons. Blue line represents light-on. (C-E) Traces of LHb neuronal calcium signals induced by astrocyte-cOpn5 before (left) and after (middle) application of various receptor antagonists. Right: bar graph showing effects of ACSF (control, C), SCH58261 (adenosine A2_A_R antagonist, D) and APV/NBQX (iGluR antagonists, E) on LHb neuronal calcium signals induced by astrocyte-cOpn5. Each circle represents one neuron.

**Figure S11.**
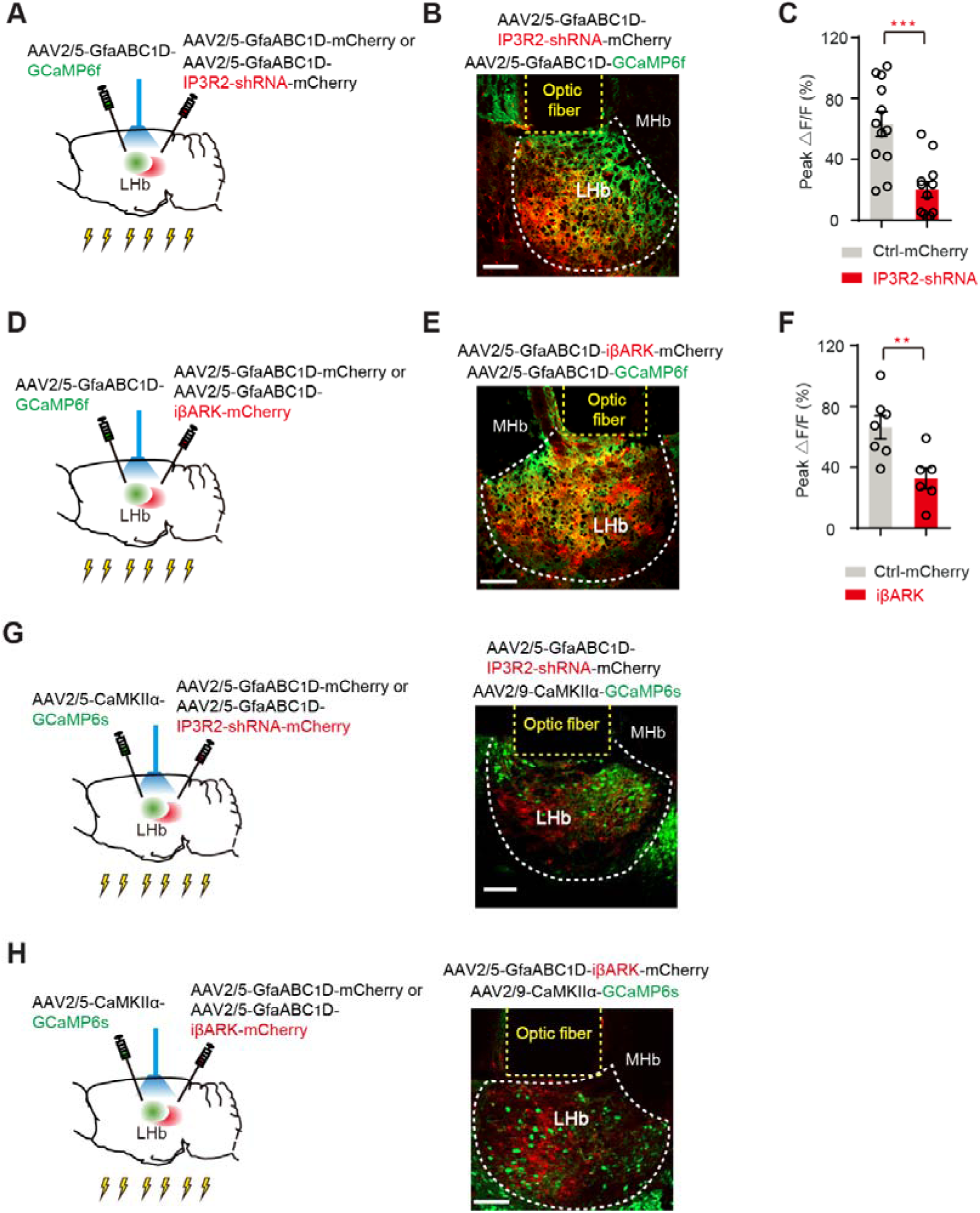
Inhibition of Astrocytic Calcium Signaling in LHb, Related to Figure 5 (A-C) Schematic (A) illustrating the viral expression of astrocyte-specific IP3R2-shRNA or Ctrl-mCherry and GCaMP6f, and fiber photometry recording of astrocytes in LHb in response to FS. Representative image (B) illustrating viral expression of IP3R2-shRNA and GCaMP6f in LHb astrocytes. Scale bar, 100 μm. Quantification of peak amplitude of FS-evoked LHb astrocytic calcium signals in mice expressing Ctrl-shRNA (grey, n = 12 mice) or IP3R2-shRNA (red, n = 12 mice) in LHb astrocytes (C). (D-F) Schematic (D) illustrating the viral expression of astrocyte-specific iβARK or Ctrl-mCherry and GCaMP6f, and fiber photometry recording of astrocytes in LHb in response to FS. Representative image (E) illustrating viral expression of iβARK and GCaMP6f in LHb astrocytes. Scale bar, 100 μm. Quantification of peak amplitude of FS-evoked LHb astrocytic calcium signals in mice expressing mCherry (grey, n = 7 mice) or iβARK (red, n = 6 mice) in LHb astrocytes (F). (G) Schematic (left) illustrating the viral expression of astrocyte-specific IP3R2-shRNA or Ctrl-mCherry and neuronal GCaMP6s, and fiber photometry recording of neurons in LHb in response to FS. Representative image (right) illustrating viral expression of IP3R2-shRNA in LHb astrocytes and GCaMP6s in LHb neurons. Scale bar, 100 μm. (H) Schematic (left) illustrating the viral expression of astrocyte-specific iβARK or Ctrl-mCherry and neuronal GCaMP6s, and fiber photometry recording of neurons in LHb in response to FS. Representative image (right) illustrating viral expression of iβARK in LHb astrocytes and GCaMP6s in LHb neurons. Scale bar, 100 μm. *P < 0.05; **P < 0.01; ***P < 0.001; ****P < 0.0001; n.s., not significant. Data are represented as mean ± SEM.

**Supplementary Table S1.**
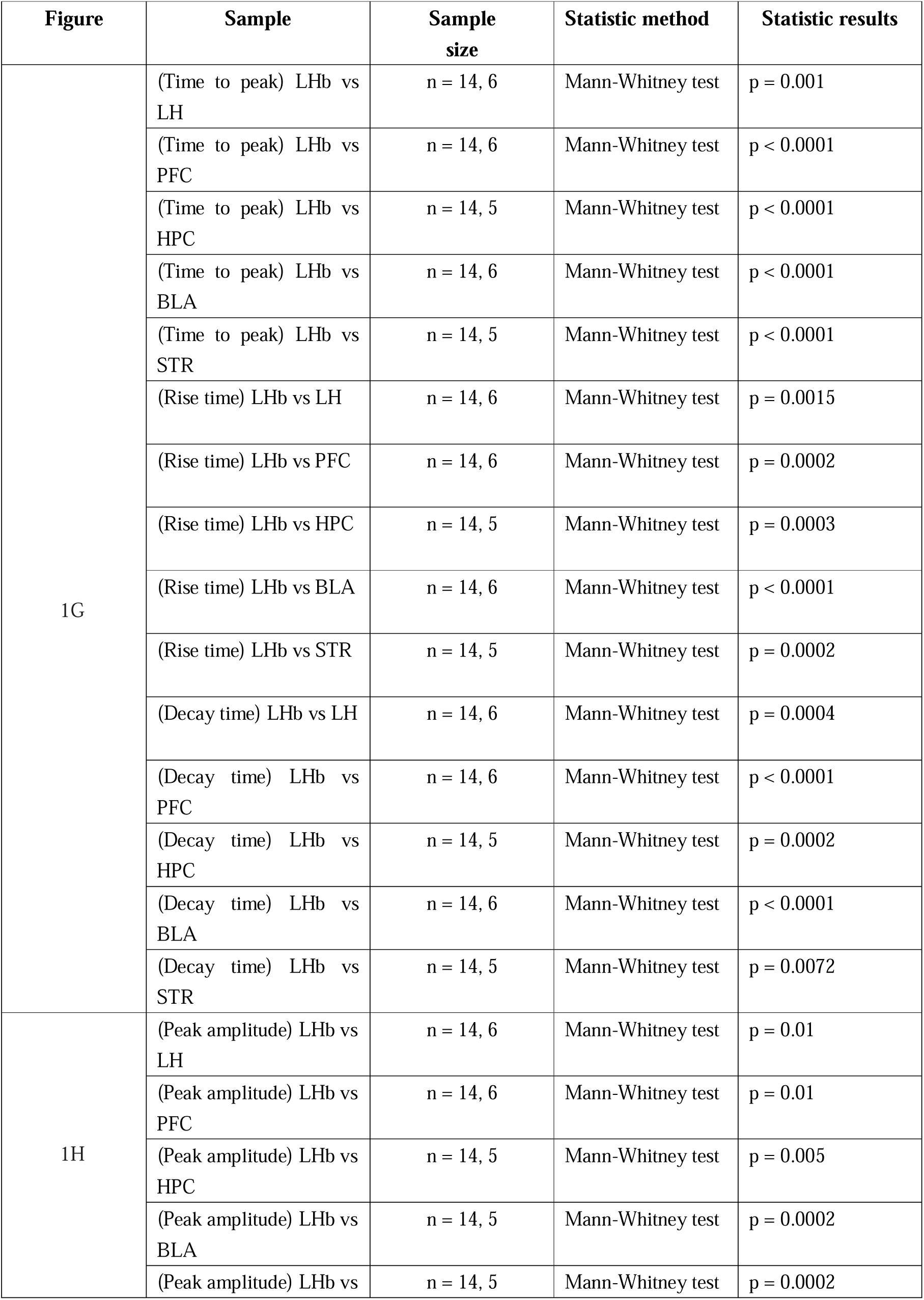
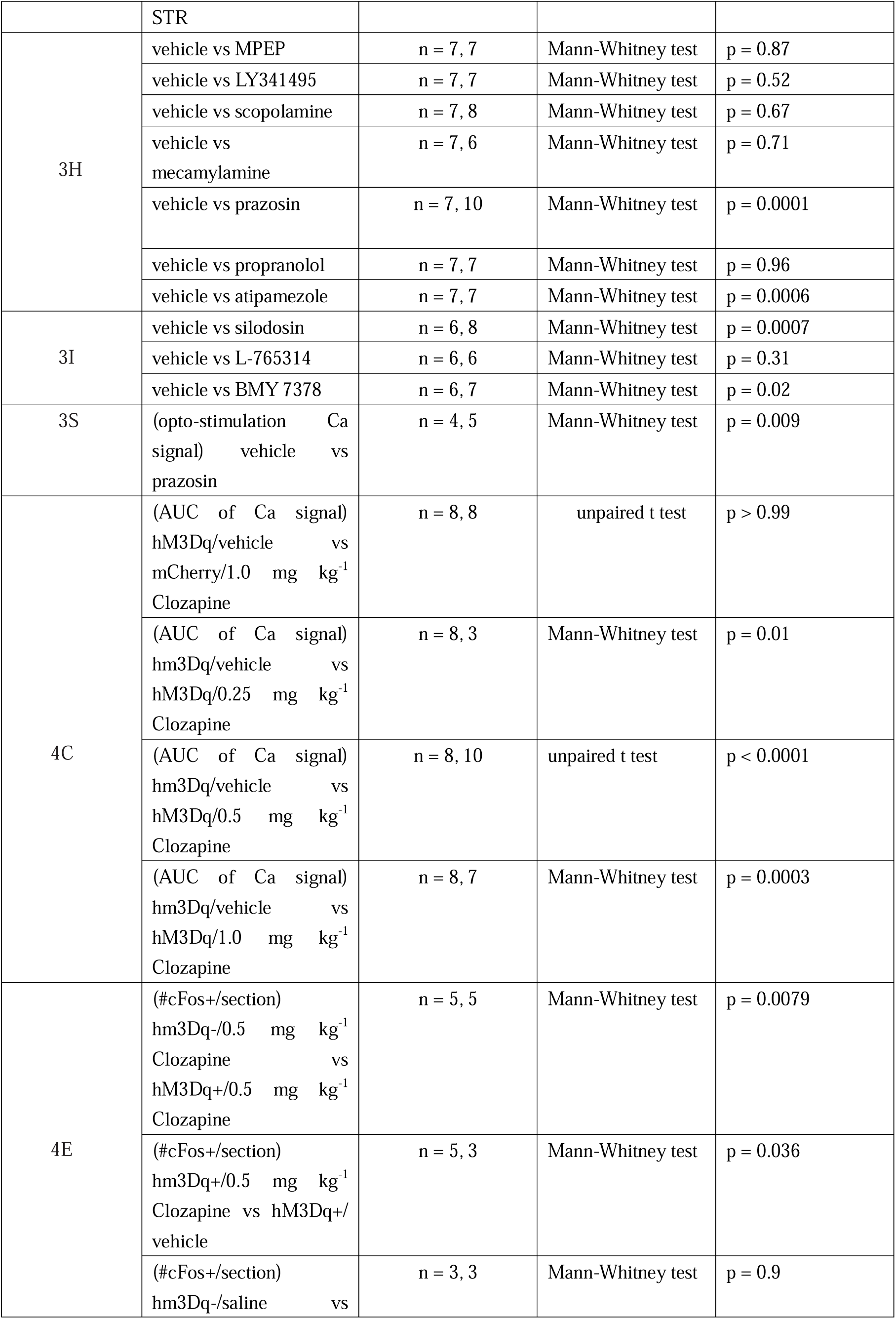
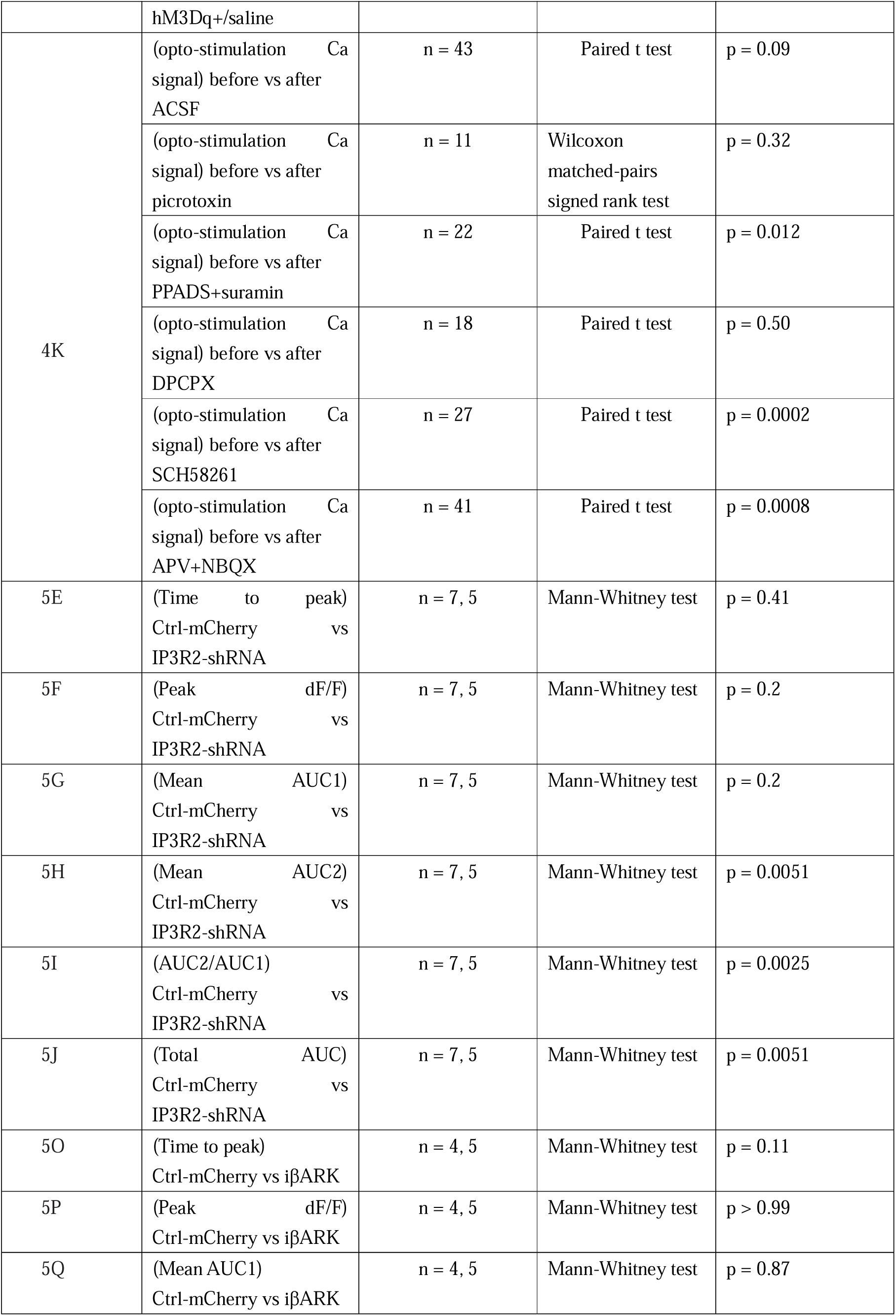
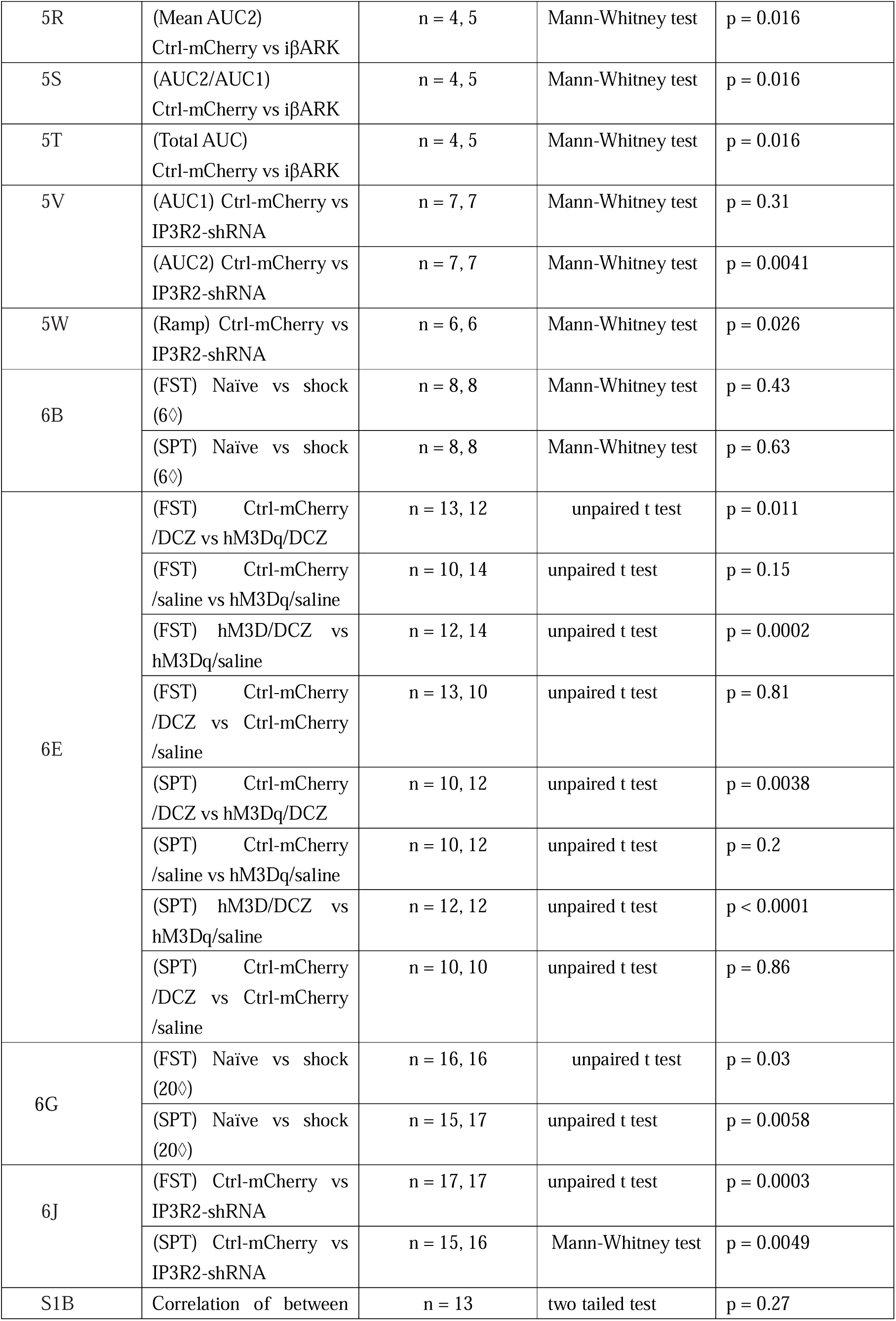
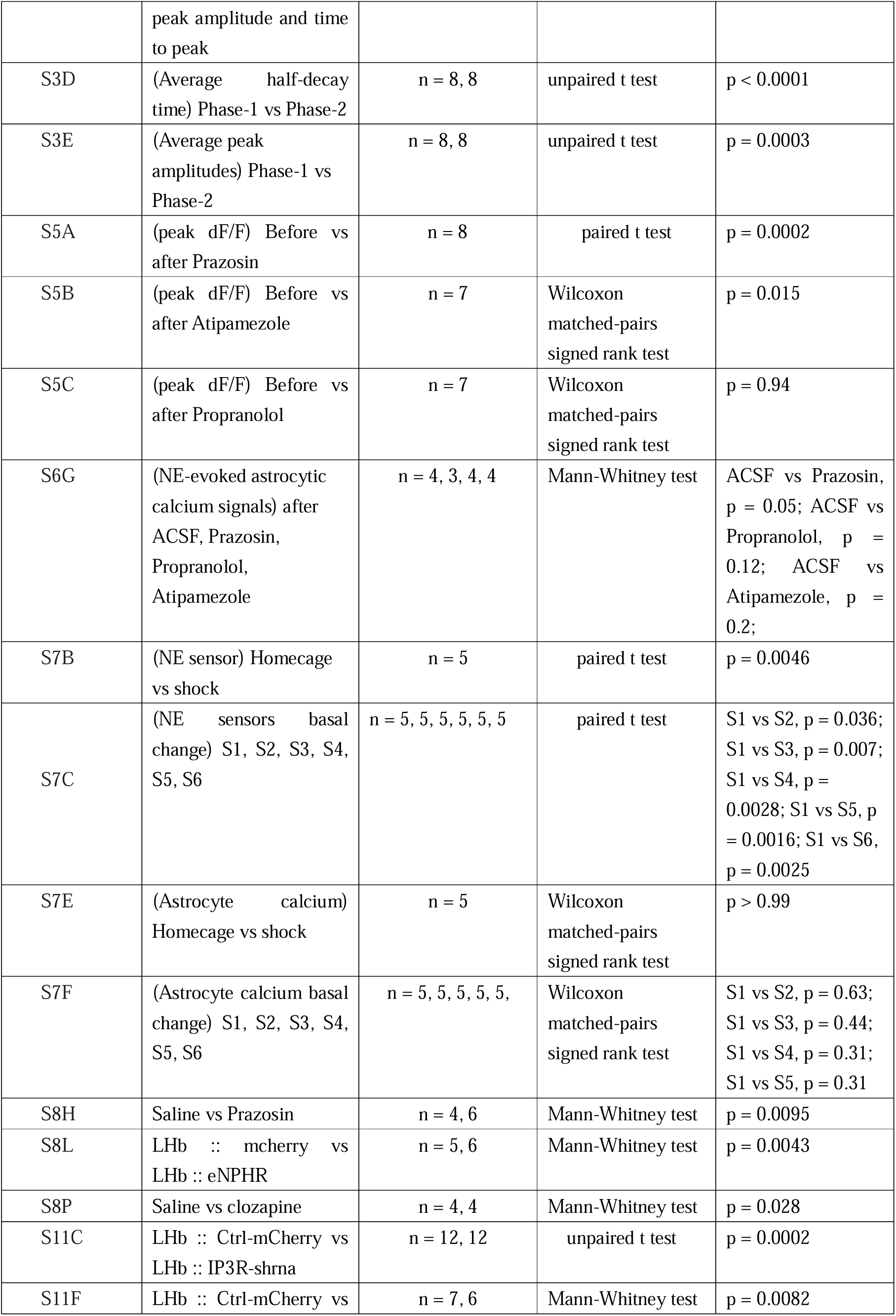
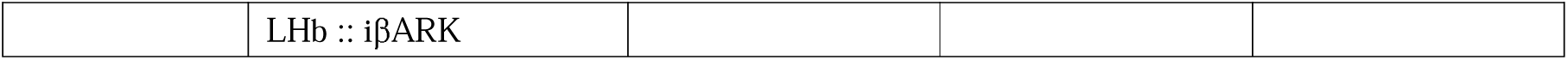
Detailed statistical information.

